# Prime-Editing in *Marchantia paleacea*: Expanding the Genome-Editing Toolbox in Bryophytes

**DOI:** 10.64898/2026.08.07.743462

**Authors:** Benoit Danilo, Aurélie Quillien, Cristina Rojas-Latorre, Zaïd Nibani, Clara Mestre, Pierre-Marc Delaux, Dominique Lauressergues, Julie Neveu

## Abstract

Since the development of CRISPR-based genome editing tools, a number of novel technologies have emerged. This includes Prime-Editing that acts as a *search and replace* genome editing tool. Prime-Editing has been deployed across multiple clades, including in a few flowering plants. Here, we report on the development of an efficient Prime Editor (PE) for the model bryophyte Marchantia. Initial tests were conducted on *Acetolactate Synthase* as a target and revealed an average efficiency above 40%. The system has been developed in the GoldenGate cloning system, facilitating construct design. The development of PE in Marchantia expands the Genome-Editing tools available for this emerging model in plant biology.

## Introduction

The next-generation genome editing technology, prime editing, can be described as a “search-and-replace” tool that precisely introduces desired insertion, deletion and DNA substitutions without generating double-strand breaks (Anzalone *et al*., 2019). The Prime Editor (PE) protein consists of an optimized *Moloney-murine leukaemia* virus reverse transcriptase (MMLV RT) fused to a modified *Streptococcus pyogenes* Cas9 nickase (Sp nCas9 H840A), which introduces a single-strand nick at the target site (Jinek *et al*., 2012). This protein complex is associated with a prime editing guide RNA (pegRNA), which includes i) a spacer sequence to guide the Cas 9 variant to the target genomic site, ii) the Primer-Binding Site (PBS) that serves as an anchoring sequence for the reverse transcriptase, and the Reverse Transcriptase Template (RTT) that encodes the desired genetic modification. Originally developed in mammalian cells, prime editing has since been successfully adapted to several models and crop plants including *A. thaliana, S. lycopersicum, S. turberosum, P. patens, O. sativa, T. aestivum*, or *Z. mays* (Vu *et al*., 2024, Perroud *et al*., 2022, Butt *et al*., 2020; Lu *et al*., 2021, Lin *et al*., 2020, Jiang *et al*., 2020). Prime editing generally exhibits a lower editing efficiency compared to the CRISPR/Cas9 system in plants. The success rate of prime editing in plant varies greatly between species, ranging from only a few percent of efficiency in *A. thaliana* to 30% in some *O. sativa* experiments (Vats *et al*., 2024). Since the original description of PE in 2019, several optimizations of the PE complex and the pegRNA have been implemented to further improve efficiency and achieve current rates. For instance, point mutations (D200N, T306L, W313F and T330P) in MMLV RT and the removal of the ribonuclease H (RNase H) domain have greatly improved prime editing efficiency from 2.1% to 11.3% in rice (Zong *et al*., 2022). Similarly in *S. lycopersicum* and in *A. thaliana*, fusion of RNA chaperones or nucleocapsid proteins to the PE, as well as overexpression of the pegRNA further increased the prime editing efficiency (Vu *et a*l., 2024). Besides flowering plants, genome editing tools have been implemented with success in Bryophytes. It includes CRISPR/Cas9, Base Editor and PE in *P. patens* (Perroud *et al*. 2022), as well as TALENs or CRISPR/Cas9 in both *M. polymorpha* and *M. paleacea* (Holman *et al*., 2026; Sugano *et al*., 2018, Rich *et al*., 2021). To this date, however, prime editing has not yet been demonstrated in *Marchantia* (Holman et al., 2026). Here, we demonstrate that prime editing can be efficiently applied to generate precise DNA modifications in *M. paleacea*. Using the enhanced plant prime editor (ePPE), we established prime editing as an efficient and precise genome editing strategy for this bryophyte species. AcetoLactate Synthase (*ALS)* has been commonly used as a reporter for genome editing in several plant species where substitutions of specific amino acid residues in *ALS* confer resistance to sulfonylurea herbicides, thereby enabling the selection of edited plants. This strategy has been implemented in maize through the introduction of the P165S, W542L, or S621I substitutions, in tomato through the introduction of the P186S, W563L or S642I substitution and in rice using the W548M substitution (Jiang *et al*., 2020; Lu *et al*., 2021, 2025). More recently, Casey and collaborators identified the P197L substitution and demonstrated that it confers resistance to chlorsulfuron in *M*.*polymorpha* when expressed as a transgene (Casey *et al*., 2023). As a proof of concept, we aimed at introducing the P197L mutation into *M. paleacea ALS* coding sequence by prime editing.

## Materials and methods

### PE editor and pegRNA design and cloning

P197L *ALS* transgene is composed of the CaMV35S promoter, the synthesized coding sequence of *ALS* and NOS terminator. The ePPE contains an engineered nCas9 (H840A), the deltaRNAseH MMLV_RT with point mutations (D200N, T306L, W313F and T330P), and a nucleocapside sequence between the nCas9 and the RT (Zong *et a*l., 2022). SV40 NLS sequences were added in N terminal and C terminal. PE promoter and terminator are MpEF1alpha and CaMV35S respectively. As selection marker, a cassette pNOS-Hygromycin-T35S was used. All the binary vectors used in this study were generated following a Golden Gate protocol (Engler *et al*., 2008). DNA sequences for level 0 were domesticated in order to remove the BsmBI, BsaI and BpiI restriction sites and cloned in the pUPD2 vector. The pegRNA was designed using the CRISPOR-Tefor website for designing the gRNA, while the RT template and PBS were designed using the PlantPegDesigner (Lin *et al*., 2021). pegRNA were synthesized by GeneArt Thermofisher. An evopreQ1 sequence was added to the pegRNA after the RT template. Two U6 endogenous *M. paleacea* promoter were used to drive the pegRNA. All these constructs are listed in the supplementary data Table 1.

### *M. paleacea* transformation

*M. paleacea* transformation was conducted as previously described (Rich *et al*. 2021). Thalli were grown *in vitro* from sterile gemmae on ½ strength Gamborg B5 (G5768, Sigma) medium supplemented with 1.4% agar (1330, Euromedex) for 6 to 8 weeks in a 16/8h photoperiod at 22 °C/20 °C. For transformation, 25 day old gemmalings were blended for 20 seconds in a sterile 250 mL stainless steel bowl (Waring, USA) in 10 mL of OM51C medium [KNO_3_ (2 mg/L), NH_4_NO_3_ (0.4 mg/L), MgSO_4_·7H_2_O (0.37 mg/L), CaCl_2_·2H_2_O (0.3 mg/L), KH_2_PO_4_ (0.275 mg/L), casamino acids (1 mg/L), Na_2_MoO_4_·2H_2_O (0.25 mg/L), CuSO_4_·5H_2_O (0.025 mg/L), CoCl_2_·6H_2_O (0.025 mg/L), ZnSO_4_·7H_2_O (2 mg/L), MnSO_4_·H_2_O (10 mg/L), H_3_BO_3_ (3 mg/L), KI (0.75 mg/L), EDTA ferric sodium (36.7 mg/L), myo-inositol (100 mg/L), nicotinic acid (1 mg/L), pyridoxine HCl (1 mg/L), thiamine HCl (10 mg/L)]. The blended plant tissues were transferred to 250 mL Erlenmeyer flasks containing 15 mL of OM51C and kept at 22 °C/20 °C at 200 RPM for 4 days. Co-cultures with *A tumefaciens* (GV3101) containing constructs were initiated at day 2, by adding 100 μL of bacterial liquid culture, acetosyringone (100 μM final) and L-glutamine (0.3 mg/L). After 3 days, plant fragments were washed 3 times with water, and plated on ½ Gamborg containing 100mg/L Cefotaxime and 10 mg/L hygromycin.

### Edited *M. paleacea* selection

Four weeks after transgenic thalli selection on 1/2 Gamborg containing 100 mg/L Cefotaxime and 10 mg/L hygromycine the regenerating thalli were transferred to 1/2 Gamborg supplemented with 150 nM chlorsulfuron to select edition events.

### Genotyping and sequencing

All regenerating thalli on chlorsulfuron media were genotyped by PCR using a couple of primer (supplementary data Table 2) flanking the desired *ALS*^*P197L*^ mutation. The PCR products were sequenced by Sanger.

## Results and Discussion

In the present study, we used the ePPE configuration (Fig. 1A) and two different pegRNA that leads to one or two nucleotide substitutions in *ALS* coding sequence (pegRNA 1 CCC > CTC and pegRNA 2 CCC > CTA) to generate the P197L modification (Fig. 1B). DNA modification of the ALS gene in *M. paleacea* induces the resistance to chlorsulfuron herbicide.We observed that one month after plant transfer on chlorsulfuron, wild-type transferred *M. paleacea* presents a yellowish color and a small size compare to the transgenic plants expressing the P197L *ALS* transgene (positive control), which were able to regenerate a proper thallus with a green color (Fig. 1C). Interestingly, an average of 40% and 45% of transgenic *M. paleacea* thalli, expressing ePPE and the pegRNA 1 and 2 respectively, presented a resistance to chlorsulfuron (Fig. 1C, F). However, we observed that transgenic thalli produced either entirely green thalli after four weeks of growth on chlorosulfuron-containing medium or only newly emerging apices corresponding to new developing thalli (Fig. 1D). These observations suggest the presence of varying degrees of mosaicism depending on the transformation event. Nevertheless, in both cases, thalli developing under selection pressure exhibited no detectable mosaicism. In the absence of selection pressure or of a phenotype associated with prime editing, however, it would still be possible to isolate non-mosaic individuals in the subsequent generation through the use of gemmae.

**Figure 1.**
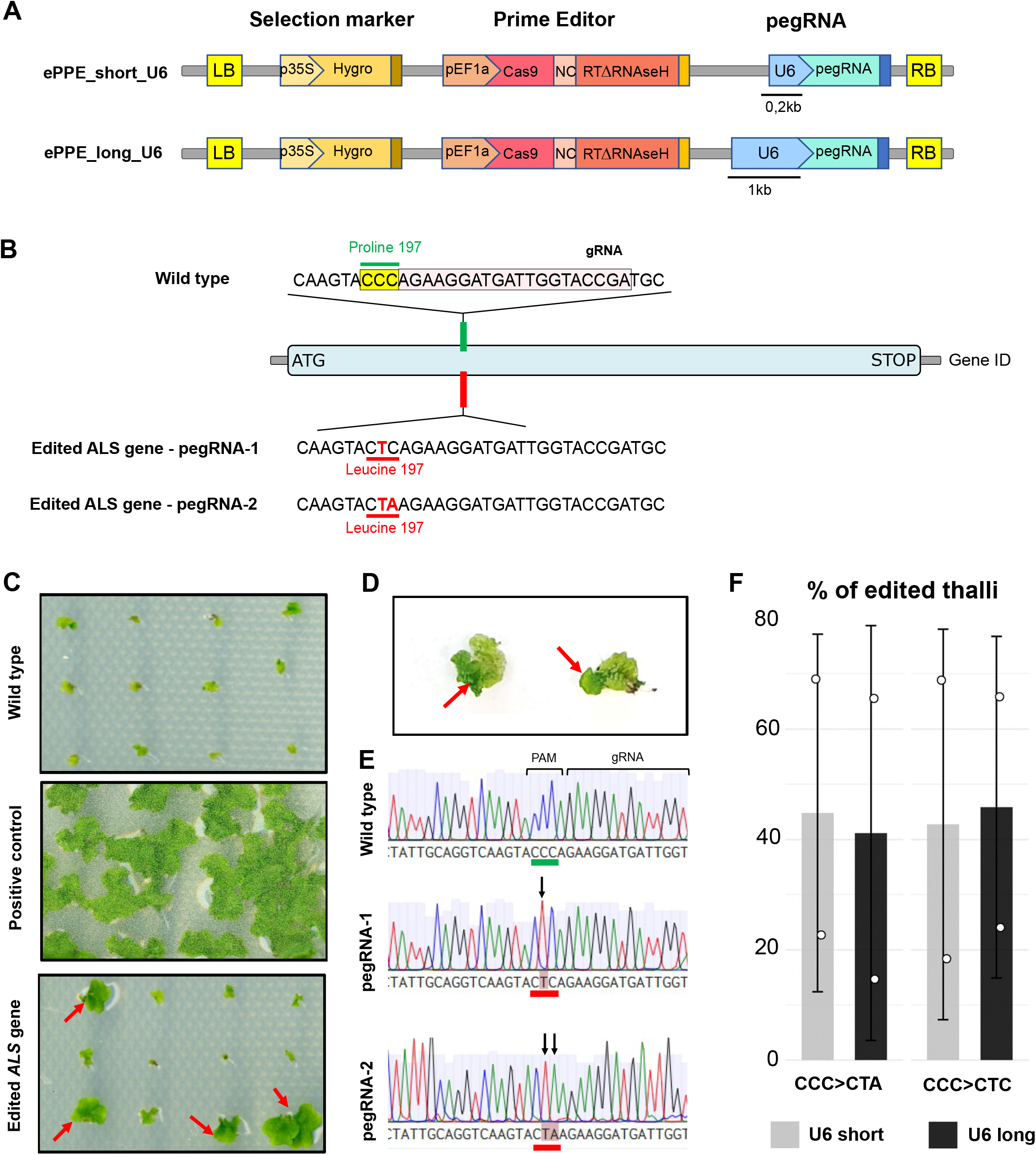
Prime-Editing in *M. Paleacea ALS* gene lead to chlorsulfuron resistance. **A**. Schematic representation of the T-DNA region of the ePPE_long_U6 and ePPE_short_U6 constructs used for Agrobacterium-mediated transformation of *M. paleacea*. Two U6 promoters were used, U6_long and U6_short, with a size of 1.9kb and 0.2kb respectively. **B**. Schematic representation of the target region in the wild-type (WT) *ALS* and *ALS* prime edited gene. The Proline 197 codon is highlighted in green. The PAM sequence boxed in yellow and guideRNA sequence is boxed in light pink. Red letters indicated modified nucleotides and the Leucine 197 is highlighted in red. The two pegRNA, pegRNA-1 and pegRNA-2, are represented with their DNA change at the ALS target. **C**. *M. paleacea* T0 regeneration thalli. The Wild type is corresponding to the unmodified *ALS* gene (Pro197). The positive control *consists of plant expressing the ALS* transgene carrying the modification P197L driven by a 35S promoter. Both transformations identified as Edited *ALS* gene-pegRNA-1 and Edited *ALS* gene-pegRNA-2 were made with the construct ePPE_short_U6. The resistant regenerating *M. paleacea* to chlorsulfuron are identified with a red arrow. **D**. Newly emerging edited apices. **E**. Chromatograms in the targeted *ALS* region of the WT and regenerated thalli with the two desired modifications. The modifications are highlighted with a black arrow and a pale red box. **F**. Prime Editing efficiency based on the average percentage of regenerating *M. paleacea* in presence of the herbicide.

To further test the intended editing, the regenerating *M. paleacea* thalli were sequenced by Sanger and both substitutions (CCC to CTC, and CCC to CTA) were correctly detected in the resistant thalli (Fig. 1E)

Additionally, two fragment of the endogenous U6 promoter, one of 1.9kb and one of 0.2kb, known to drive effective gRNA expression in the context of CRISPR/Cas9, were also tested. The average percentage of edited *M. palaecea* was not statistically different between the two tested versions of the MpaU6_pro_ (Fig. 1F). Altogether, this data suggest that prime editing can be carried out successfully in *M. paleacea*. More than 40% of editing efficiency can be reached with either one or two nucleotide substitutions, as well as with both the long and short version of the U6 promoter.

This promising result offer new perspective for genetic manipulation of *M. paleacea*. To our knowledge this is the first example of Prime Editing in the emerging model genus *Marchantia*. This new example of application of last generation genome editing tools expands the genome editing toolbox for Bryophytes. Recently, other variants of PE have been developed for prime editing in mammalian cells such as PEmax, PEmax** or PE7 (Chen *et al*., 2021; Liu *et al*., 2025). These new variants were applied to performed plant prime editing in tomato, rice and *A. thaliana* (Jiang *et al*., 2022; Vu *et al*., 2024) and have a better efficiency compared to the ePPE configuration. It the future, it will be interesting to test these new variants in *M. paleacea* with the objective to improve the efficiency of Prime Editing. Other strategies were developed such as the use of new reverse transcriptase evoTf1 that improve prime editing in mammalian cells, and in tomato calli with an average of 16% efficiency (Doman *et al*., 2023; Van Vu *et al*., 3 2026).

Expanding the genome editing toolbox for Bryophytes with this simple example of Prime Editing will facilitate functional analysis of this understudied plant clade, with direct impact on our understanding of plant evolution and the dissection of bryophyte-specific mechanisms and traits.

## Acknowledgments

The authors thank the TYPEX consortium, in particular Fabien Nogué, for initial discussions and insightful comments on the PE design. This work was supported by the “Laboratoires d’Excellence (LABEX)” TULIP (ANR-10-LABX-41)” and by the “École Universitaire de Recherche (EUR)” TULIP-GS (ANR-18-EURE-0019). This project has received funding from the European Research Council (ERC) under the European Union’s Horizon 2020 research and innovation program (grant agreement No 101001675 - ORIGINS) to P-M.D. This work is part of the TYPEX project of the Sélection Végétale Avancée research program and benefited from government funding managed by the National Research Agency under the France 2030 program, reference ANR-22-PESV-0002.

