## Supplementary data for "Prime-Editing in *Marchantia paleacea*: Expanding the Genome-Editing Toolbox in Bryophytes"

Table 1 : List of plasmids used for Prime Editing of *MarpalALS* gene in this study

| Plasmids ID | Plasmid description |
|---|---|
| 2_TYPEX_001 | p35S::HygromycinR-Tnos -- pMpaEF1a::ePPE-T35S -- pMpaU6-1short::MpaALS_pegRNA2 CCC->CTA |
| 2_TYPEX_003 | p35S::HygromycinR-Tnos -- pMpaEF1a::ePPE-T35S -- pMpaU6-long::MpaALS_pegRNA2 CCC->CTA |
| 2_TYPEX_005 | p35S::HygromycinR-Tnos -- pMpaEF1a::ePPE-T35S -- pMpaU6-1short::MpaALS_pegRNA1 CCC->CTC |
| 2_TYPEX_007 | p35S::HygromycinR-Tnos -- pMpaEF1a::ePPE-T35S -- pMpaU6-long::MpaALS_pegRNA1 CCC->CTC |

Table 2 : List of primers used for *MarpalALS* gene genotyping in this study

| Primers ID | Primer sequence |
|---|---|
| 2436 | CCACCAGGCTCTTACACGTT |
| 2437 | AGCTTCCCTGATTACGCGAG |

>2_TYPEX_001
CTGGTGGCAGGATATATTGTGGTGTAAACAAATTGACGCTTAGACAACTTAATAACACATTGCGGACG
TTTTTAATGTACTGGGGTTGAACACTCTGtgccgaattcggatccagcgtcgatctagtaacatagat
gacaccgcgcgcgataatttatcctagtttgcgcgctatattttgttttctatcgcgtattaaatgta
taattgcgggactctaatcataaaaacccatctcataaataacgtcatgcattacatgttaattatta
catgcttaacgtaattcaacagaaattatatgataatcatcgcaagaccggcaacaggattcaatctt
aagaaactttattgccaaatgtttgaacgatctgcttgacaagcctattcctttgccctcggacgagt
gctggggcgtcggtttccactatcggcgagtacttctacacagccatcggtccagacggccgcgcttc
tgcgggcgatttgtgtacgcccgacagtcccggctccggatcggacgattgcgtcgcatcgaccctgc
gcccaagctgcatcatcgaaattgccgtcaaccaagctctgatagagttggtcaagaccaatgcggag
catatacgcccggagccgcggcgatcctgcaagctccggatgcctccgctcgaagtagcgcgtctgct
gctccatacaagccaaccacggcctccagaagaagatgttggcgacctcgtattgggaatccccgaac
atcgcctcgctccagtcaatgaccgctgttatgcggccattgtccgtcaggacattgttggagccgaa
atccgcgtgcacgagatgccggacttcggggcagtcctcggcccaaagcatcagctcatcgagagcct
gcgcgacggacgcactgacggtgtcgtccatcacagtttgccagtgatacacatggggatcagcaatc
gcgcatatgaaatcacgccatgtagtgtattgaccgattccttgcggtccgaatgggccgaacccgct
cgtctggctaagatcggccgcagcgatcgcatccatggcctccgcgaccggctgcagttatcatcatc
atcatagacacacgaaataaagtaatcagattatcagttaaagctatgtaatatttacaccataacca
atcaattaaaaaatagatcagtttaaagaaagatcaaagctcaaaaaaataaaaagagaaaagggtcc
taaccaagaaaatgaaggagaaaaactagaaatttacctgcagaacagcgggcagttcggtttcaggc
aggtcttgcaacgtgacaccctgggcacggcgggagatgcaataggtcaggctctcgctgaattcccc
aatgtcaagcacttccggaatcgggagcgcggccgatgcaaagtgccgataaacataacgatctttgt
agaaaccatcggcgcagctatttacccgcaggacatatccacgccctcctacatcgaagctgaaagca
cgagattcttcgccctccgagagctgcatcaggtcggacacgctgtcgaacttttcgatcagaaactt
ctcgacagacgtcgcggtgagttcaggctttttcattgcgtgtcctctccaaatgaaatgaacttcct
tatatagaggaagggtcttgcgaaggatagtgggattgtgcgtcatcccttacgtcagtggagatgtc
acatcaatccacttgctttgtagacgtggttggaacctcttctttttccacgatgctcctcgtgggtg
ggggtccatctttgggaccactgtcggcagagagatcttgaatgatagcctttcctttatcgcaatga
tggcatttgtaggagccaccttccttttctactgtcctttcgatgaagtgacagatagctgggcaatg
gaatccgaggaggtttcccgaaattatcctttgttgaaaagtctcaatagccctttgatcttctgaga

ctgtatctttgacatttttggagtagaccagagtgtcgtgctccaccatgttgacctccGCAAGAATT
CAAGCTTAGCGatctggattttagtactggattttggttttaggaattagaaattttattgatagaag
tattttacaaatacaaatacatactaagggtttcttatatgctcaacacatgagcgaaaccctatagg
aaccctaattcccttatctgggaactactcacacattattatggagaaactcgagcttgtcgatcgac
tctagctagagaagcCTAGAGCGGTTGATCTGTGAGGTCTGGCCTTGTGCCATGCGCTTCCGCGAGAA
TATCCAGACAGTTGTGTTGGAGGCCCTCCTCTGGGAGTGGCAGCAGTGTCGCCGGATTGAGGGCGACC
ACTGGGCCGAACTGGACGCGATCGGTGTCGAGGAGGAGCGCTTGGTAATGGGTCATGCGCGCATTGGA
GAGCCACCTATCCGGCGGCTGTTTGACGAGGGCTTCCACGGCATGTGGCGCGAGGATCACCAGCGGCT
GGCCCATTGTGAGTTTGCCGGCATCCTTTGTGAGCACGGCAATCGCGGCGACCATGCGCAGACACGGT
GGCCAGCCGGCGGCCACTGGATCGAGTTTTTGGAGAGATAGGCCACTGGCCTCCTCCACGGGCCGAG
CTTCTGTGTGAGCACGCCCTTCGCGTAGCCTTGCTTCTCATCGACGAACAGCTCGAACGGCTTTGTGA
GATCTGGGAGCCCGAGCGCTGGGGCGGTGAGGAGCGCTTGTTTGATCTCTTGGTAGGCTTTCTGCTGA
TCCGGCCCCCAGTTAAACAGTGTGCCCGGCTTGGTGAGTGGGTAGAGTGGCGCGGCCATCTCGGCAAA
GCCCGGGATGAAGAGGCGACAGAAGCCGGCTTTGCCGAGAAACTCGCGCAGCTGGCGTGGTGTTTTCG
GTGTCGGTTGCCCCATCACTGTTTCTTTGCGCGCTTCGGTGAGCCAGCGTTGGCCTTCCTTGAGCAGA
TACCCGAGGTACTTCACTTGCTTCTGGCAGATTTGGGCCTTTTTCGCGGACGCGCGATAGCCGAGGTT
CCCGAGTGTTTGGAGGAGCGCGCGTGTGCCTTGTTGACAATCCAGCTCGGAGGTCGCGGCGAGGAGGA
GATCATCCACATACTGGAGGAGGATGAGGTCCGGATGCTGGATGCGAAAATCCGCGAGGTCCCTATGG
AGCGCCTCGTTGAAGAGTGTCGGGCTGTTCTTGAAGCCTTGCGGGAGGCGTGTCCATGTCAGTTGGCC
GGAGATGCCCATCTCCGGGTCGCGCCACTCGAAGGCGAAGAGCGGCTGGCTTGTTGGGTGGAGGCGGA
GACAGAAGAAGGCGTCTTTGAGGTCCAGCACTGTATACCACTGGTGGGATGGTGGGAGCCCGGAGAGC
AGATTATACGGGTTCGGGACGGTCGGGTGGATaTCTTCCACGCGCTTGTTGACCTCGCGGAGATCTTG
GACCGGGCGATAGTCGTTGGTGCCCGGCTTTTTGACCGGGAGGAGTGGTGTATTCCACGGGGATTGGC
ACGGCACGAGAATGCCTTGATCGAGGAGGCGCTGAATGTGCGGCTTAATCCCGAGCCTCGCCTCTTGG
CTCATCGGGTACTGCTTGATGGACACCGGTGTGGAGGTCGCCTTCAGCGGGATAATGAGTGGGGCTTG
GCGGACGGCGAGGCCCATGCCGCCGGTTTCGGCCCACGCTTGTGGGAAGTCGCTGAGCCATGTGCTGC
CGAGGGACACGTCCGGTTCTTTGGAGGTTTCGTGGAGGCGATACTCATCCTCGATGTTGAGGGTAGGC
CTAGATCCTCCGGAAGAGCCTCCGCTAGATTCTGGTGTAGCGCTTTCGGATGTTCCTGGTGTCTCGGA
GCCGCTAGATCCTCCGGAAGATCCTCCGCTGAGCTCCACCTTCCGCTTCTTCTTTGGCGAACCAAGGA
GGGATGTTTGTGGCCTTGGGCCGCGTGGGCCGCGCGGCTTTTTCGGGCAGTCTTTGGCCCAGTGGCCC
TTCTCCTTGCAGTAGGCACACTGATCCCTATCGAGCTGGGACCTTCTGCGTTCTCCGCCCTGGCGGTC
CTGCTTTTGGCCGGACACCACTGTGGCAGATTCTGGGGTGGCCGACTCGGAGGTGCCTGGCGTTTCGC
TCCCGGAACCTGAATCACCACCGAGCTGTGAGAGATCGATCCTAGTCTCGTAGAGTCCAGTGATAGAC
TGATGGATGAGGGTAGCATCGAGCACTTCTTTGGTAGAGGTGTATCTCTTCCTATCGATGGTTGTATC
GAAGTACTTGAAAGCAGCAGGAGCACCGAGGTTGGTAAGGGTGAAGAGATGGATGATGTTCTCTGCCT
GTTCCCTGATAGGCTTATCTCTGTGCTTGTTGTAAGCAGACAACACCTTATCGAGGTTTGCATCAGCG
AGGATCACCCTTTTAGAGAACTCAGAGATCTGCTCGATGATCTCATCCAAGTAGTGCTTGTGCTGCTC
AACGAAAAGTTGCTTCTGCTCGTTATCTTCTGGAGATCCCTTCAACTTCTCGTAGTGAGAAGCGAGGT
AAAGAAAGTTAACGTACTTAGATGGGAGAGCAAGCTCGTTTCCCTTTTGAAGCTCACCAGCAGAAGCG
AGCATCCTCTTTCTACCGTTCTCGAGTTCGAAGAGTGAGTACTTTGGGAGCTTGATGATGAGATCCTT
CTTAACCTCTTTGTATCCCTTAGCCTCGAGGAAATCGATTGGGTTCTTCTCGAAAGATGACCTTTCCA
TGATAGTGATTCCGAGAAGTTCCTTAACAGACTTGAGCTTCTTACTCTTTCCCTTCTCAACCTTAGCC
ACAACGAGAACAGAGTAAGCCACGGTAGGAGAATCGAAACCACCGTATTTCTTAGGGTCCCAATCCTT
CTTCCTAGCAATGAGCTTATCAGAGTTCCTCTTAGGGAGGATAGACTCTTTAGAGAATCCACCGGTCT
GCACCTCGGTTTTCTTAACGATGTTCACCTGTGGCATAGAGAGCACCTTTCTAACGGTAGCGAAATCC
CTTCCCTTATCCCACACGATCTCACCTGTTTCACCGTTTGTCTCGATGAGTGGCCTCTTTCTGATCTC
ACCGTTAGCGAGGGTAATCTCGGTCTTGAAGAAATTCATGATGTTAGAGTAGAAGAAATACTTAGCGG
TAGCCTTTCCGATCTCTTGCTCAGACTTAGCGATCATCTTCCTCACATCGTACACCTTGTAATCACCG
TACACGAACTCTGACTCGAGCTTAGGATACTTCTTGATGAGAGCGGTTCCAACAACAGCGTTAAGGTA
AGCATCGTGAGCGTGGTGGTAGTTGTTGATTTCCCTCACCTTGTAGAATTGGAAATCCTTTCTGAAAT
CAGACACGAGCTTTGACTTGAGGGTGATAACCTTCACTTCCCTGATCAACTTATCGTTCTCATCGTAC
TTGGTGTTCATCCTAGAATCGAGGATCTGTGCAACGTGCTTAGTGATCTGCCTGGTTTCCACAAGCTG

CCTCTTGATGAATCCTGCCTTATCCAATTCAGAGAGTCCTCCCCTCTCAGCCTTAGTCAAGTTATCGA
ACTTTCTCTGAGTGATGAGCTTAGCGTTGAGGAGCTGCCTCCAATAGTTCTTCATTTTCTTCACAACC
TCTTCACTTGGCACGTTATCACTCTTACCCCTGTTCTTATCAGACCTGGTGAGCACCTTGTTATCGAT
AGAATCATCCTTCAAGAATGACTGTGGCACGATAGCATCAACATCGTAATCAGAGAGCCTGTTGATAT
CCAACTCTTGATCCACATACATATCCCTTCCGTTCTGGAGGTAGTAGAGGTAGAGCTTCTCATTCTGG
AGCTGAGTGTTCTCAACAGGGTGCTCTTTGAGGATCTGAGATCCAAGCTCTTTGATACCTTCCTCGAT
CCTCTTCATCCTTTCCCTAGAGTTCTTCTGTCCCTTCTGAGTGGTCTGGTTCTCTCTAGCCATTTCGA
TCACGATGTTCTCAGGCTTATGCCTTCCCATCACCTTCACCAACTCATCCACAACCTTCACAGTCTGG
AGGATTCCCTTCTTGATTGCAGGAGATCCAGCGAGGTTAGCGATATGCTCATGGAGACTATCACCCTG
TCCTGAAACCTGAGCCTTCTGGATATCCTCTTTAAAGGTGAGAGAATCATCGTGGATGAGCTGCATGA
AGTTTCTGTTAGCGAATCCATCAGACTTGAGGAAATCAAGGATTGTCTTTCCAGACTGCTTATCCCTG
ATTCCGTTAATGAGCTTTCTTGAGAGCCTTCCCCAACCAGTGTATCTTCTTCTCTTCAACTGCTTCAT
CACCTTATCATCGAAGAGATGAGCGTAGGTCTTGAGCCTTTCTTCAATCATCTCTCTATCTTCAAAGA
GGGTGAGGGTAAGAACGATATCCTCCAAGATATCCTCGTTTTCCTCGTTATCCAAGAAATCCTTATCC
TTAATGATCTTGAGGAGATCGTGGTAGGTTCCGAGAGATGCGTTGAACCTATCCTCAACACCAGAAAT
CTCAACTGAATCGAAGCACTCGATTTTCTTGAAGTAATCCTCTTTGAGCTGCTTCACGGTCACCTTTC
TGTTGGTCTTGAACAAGAGATCAACGATAGCCTTCTTTTGCTCACCTGACAAAAAAGCAGGCTTCCTC
ATTCCCTCGGTCACGTACTTAACCTTGGTCAACTCGTTGTACACGGTGAAGTACTCGTAGAGCAAAGA
GTGCTTAGGGAGCACCTTCTCGTTTGGAAGGTTCTTATCGAAGTTGGTCATCCTCTCGATGAAAGACT
GAGCACTAGCACCCTTATCCACCACCTCTTCGAAGTTCCAAGGGGTGATGGTTTCCTCAGACTTTCTG
GTCATCCAAGCGAATCTTGAGTTTCCTCTAGCGAGAGGTCCCACGTAGTAAGGGATTCTGAAGGTGAG
AATCTTCTCAATCTTTTCCCTGTTATCCTTGAGGAATGGGTAGAAATCCTCTTGCCTTCTAAGGATAG
CGTGCAACTCTCCGAGGTGGATCTGATGAGGGATAGATCCGTTATCGAAGGTCCTCTGCTTTCTGAGA
AGATCCTCTCTATTGAGCTTCACGAGGAGTTCCTCGGTTCCATCCATCTTCTCGAGGATAGGCTTGAT
GAACTTGTAGAACTCTTCTTGAGATGCACCACCATCGATGTAACCAGCGTATCCGTTCTTAGACTGAT
CGAAGAAAATCTCTTTGTACTTCTCTGGGAGCTGCTGTCTAACAAGAGCCTTGAGAAGTGTGAGATCC
TGGTGGTGCTCATCGTATCTCTTGATCATAGAAGCTGAGAGTGGAGCCTTGGTGATCTCGGTGTTCAC
TCTGAGGATATCACTGAGGAGGATAGCATCAGAGAGGTTCTTAGCAGCGAGGAACAAATCAGCGTACT
GATCTCCGATCTGAGCGAGGAGGTTATCGAGATCATCATCGTAGGTATCCTTTGAGAGCTGGAGCTTT
GCATCCTCAGCGAGATCGAAGTTAGACTTGAAGTTAGGGGTGAGTCCGAGAGAGAGAGCGATCAAGTT
TCCGAAAAGTCCGTTCTTCTTCTCACCAGGGAGCTGAGCAATGAGGTTCTCAAGCCTTCTTGACTTAG
AGAGCCTAGCAGAGAGGATAGCCTTAGCATCCACACCTGAAGCGTTGATAGGGTTCTCTTCGAAAAGC
TGGTTGTAGGTCTGCACGAGCTGGATGAACAACTTATCCACATCAGAGTTATCAGGGTTGAGATCACC
CTCGATGAGGAAGTGTCCTCTGAACTTGATCATGTGAGCGAGAGCGAGGTAGATGAGCCTGAGATCAG
CCTTATCAGTAGAATCAACGAGCTTCTTTCTGAGGTGGTAGATAGTAGGGTACTTCTCGTGGTATGCC
ACCTCATCAACGATGTTTCCGAAGATAGGGTGCCTCTCGTGCTTCTTATCTTCTTCCACGAGGAATGA
CTCTTCGAGCCTGTGGAAGAATGAATCATCCACTTTAGCCATCTCGTTAGAGAAGATCTCTTGGAGGT
AGCAGATCCTGTTCTTTCTTCTGGTGTACCTTCTTCTAGCGGTTCTCTTGAGTCTGGTAGCCTCAGCA
GTTTCACCAGAATCGAAGAGGAGAGCACCGATAAGGTTTTTCTTGATAGAGTGCCTATCGGTGTTTCC
GAGAACCTTGAACTTCTTAGATGGCACCTTGTACTCATCGGTGATCACAGCCCATCCCACAGAGTTAG
TTCCGATATCGAGTCCGATAGAGTACTTCTTATCAACCTTTCTCTTCTTCTTAGGAGATTCAAATTCA
GATCCATCAGCAGTTCTCTTCATCATTcaacctttctgcaggcacatcaatactgttgaggcaacacc
catagaagcacggaaccaaggagcaccgacgtccatagcacacagccaacctcgctgaccttcgactc
ctcttcaagtcaaggatagacaaatctatcatgcaggaaacttacgaggcacctccgcaccagtgacc
gaactgaacaccttaccgagataacaggactgaccaaagtgggatgtacttggctaaaacgaacctaa
aaaatcactcgctcgaggccccaatcgacaccacgtaaaagctccagtcgggtttcaggtctagacaa
tcacaatatcgagcaatcgtaacgacgtacagtgacgagggcacaaggcgcataccagtagacaagat
gcagctcttcgatgtcgatgctctccacgcaagagagaaagaagtgaaaggaactctagggttgcggc
gatcgattttaagccagacgctccactgcctcccagcgagctttgagagcaagcgcgaccacagcact
agacgccaaagtagaagaaattagggaatgattagcgaaattgcggagggttttgatataattaaata
cctaggtctgccccgtttataataacgagcgttcgataaggactttgcatcggctccgttcaagttat
tgcgacttagggtcggcggccggccattcgcagataggcggccaattcagcatgcgagggcaagttgg

gaggccgacgacgaagggttaggaatattcaaagcctcttttccatttatatattgccctgtccatgc
cccttcggactctttgtcgtgactgccggccagcatcgtcatccgaattgctcacactcctccgtctg
tcacatcgtccgtgcttagggccggcacgccctttcaatttaacattccaaaataagatgacgtaggt
ttagcgaacccgactgctgataccacaccgattgctccggtttttcgagtaccttttccgcgcttatt
acaagctgcgagcaaactttctagaaagtagagaggtggccgaagtcgcttttggcctcccgcgcctt
tctccttctgcacatacttccaaacggtggactagcatcccatagatccatgacatgcacgcaatcac
atgagattttactgttgctttcttcttctggcctcaccgtcctccccgtcctcccttcgcttttggg
acgcgggaaaacctagaaggcttctgattgaagtttccttttctgcttgtcggcctttacactttgga
gaacgaagttctaatctctcttgaagttgaccaaatattagtatagtgaactgatcgagaagatgcgg
gtccgatccaataggagagaacaaaatattacgagacagaaagagataattaagcaatttatatgcac
tttaaaatggaatcttttcgattatcatcaatgagcattatattgtgaaacggatttaaaggattagc
tattttgaaaggtttttttaatctgtttttttaagatctttagcaaattctttttaagttacccaaata
tcgatacccgttaacaatcaaaaatcgatgacagagttgtgaatcatcatcaagatctgcaaatttga
tatgattttgaaaatttgagatacaaatattgttaaaataatgcatttaactgggaaatctatcaact
taccttacaattgagttattttgatatgtctctcaacaatgtgtgtgactcatttgctccactagaat
tcgagctcagcgaaaaaaagcaccgactcggtgccacttttttcaagttgataacggactagccttatt
ttaacttgctatttctagctctaaaacagacataaaaaacaaaaaaaTTTCTAGTTGGTTTAACGCGT
AACTAGATAGAACCGCGTCAAGAGAGAGAGGTACCAATCATCCTTCTTAGTACTTGACGCACCGACTC
GGTGCCACTTTTTCAAGTTGATAACGGACTAGCCTTATTTTAACTTGCTATTTCTAGCTCTAAAACAG
AAGGATGATTGGTACCGAcTATGgcaaggtgcaatttacggatttatacactacatttgcattgtgcg
ttttgcttatagttttactctcgataagcaaaccacgtggtcccttctccgtcattgctgtcgtgaat
ttctgaagtaaccggaaaatgtttcttggccgtcatgagttcaatcctgctgtgcgatcgcgttctgg
ctcggtccaataccactccgtctgctcgtttcggtatatatatcggacctagaggagaatggagatcg
tcaagacatcccaccatgtccacTTACGAGGATGCACATGTGACCGAGGGACACGAAGTGATCCGTTT
AAACTATCAGTGTTTGACAGGATATATTGGCGGGTAAACCTAAGAGAAAAGAGCGTTTATTAGAATAA
TCGGATATTTAAAAGGGCGTGAAAAGGTTTATCCGTTCGTCCATTTGTATGTGCATGCCAACCACAGG
GTTCCCCTCGGGAGTCAGCCGTGCGGCTGCATGAAATCCTGGCCGGTTTGTCTGATGCCAAGCTGGCG
GCCTGGCCGGCCAGCTTGGCCGCTGAAGAAACCGAGCGCCGCCGTCTAAAAAGGTGATGTGTATTTGA
GTAAAACAGCTTGCGTCATGCGGTCGCTGCGTATATGATGCGATGAGTAAATAAACAAATACGCAAGG
GGAACGCATGAAGGTTATCGCTGTACTTAACCAGAAAGGCGGGTCAGGCAAGACGACCATCGCAACCC
ATCTAGCCCGCGCCCTGCAACTCGCCGGGGCCGATGTTCTGTTAGTCGATTCCGATCCCCAGGGCAGT
GCCCGCGATTGGGCGGCCGTGCGGGAAGATCAACCGCTAACCGTTGTCGGCATCGACCGCCCGACGAT
TGACCGCGACGTGAAGGCCATCGGCCGGCGCGACTTCGTAGTGATCGACGGAGCGCCCCAGGCGGCGG
ACTTGGCTGTGTCCGCGATCAAGGCAGCCGACTTCGTGCTGATTCCGGTGCAGCCAAGCCCTTACGAC
ATATGGGCCACCGCCGACCTGGTGGAGCTGGTTAAGCAGCGCATTGAGGTCACGGATGGAAGGCTACA
AGCGGCCTTTGTCGTGTCGCGGGCGATCAAAGGCACGCGCATCGGCGGTGAGGTTGCCGAGGCGCTGG
CCGGGTACGAGCTGCCCATTCTTGAGTCCCGTATCACGCAGCGCGTGAGCTACCCAGGCACTGCCGCC
GCCGGCACAACCGTTCTTGAATCAGAACCCGAGGGCGACGCTGCCCGCGAGGTCCAGGCGCTGGCCGC
TGAAATTAAATCAAAACTCATTTGAGTTAATGAGGTAAAGAGAAAATGAGCAAAAGCACAAACACGCT
AAGTGCCGGCCGTCCGAGCGCACGCAGCAGCAAGGCTGCAACGTTGGCCAGCCTGGCAGACACGCCAG
CCATGAAGCGGGTCAACTTTCAGTTGCCGGCGGAGGATCACACCAAGCTGAAGATGTACGCGGTACGC
CAAGGCAAGACCATTACCGAGCTGCTATCTGAATACATCGCGCAGCTACCAGAGTAAATGAGCAAATG
AATAAATGAGTAGATGAATTTTAGCGGCTAAAGGAGGCGGCATGGAAAATCAAGAACAACCAGGCACC
GACGCCGTGGAATGCCCCATGTGTGGAGGAACGGGCGGTTGGCCAGGCGTAAGCGGCTGGGTTGTCTG
CCGGCCCTGCAATGGCACTGGAACCCCCAAGCCCGAGGAATCGGCGTGACGGTCGCAAACCATCCGGC
CCGGTACAAATCGGCGCGGCGCTGGGTGATGACCTGGTGGAGAAGTTGAAGGCCGCGCAGGCCGCCCA
GCGGCAACGCATCGAGGCAGAAGCACGCCCCGGTGAATCGTGGCAAGCGGCCGCTGATCGAATCCGCA
AAGAATCCCGGCAACCGCCGGCAGCCGGTGCGCCGTCGATTAGGAAGCCGCCCAAGGGCGACGAGCAA
CCAGATTTTTTCGTTCCGATGCTCTATGACGTGGGCACCCGCGATAGTCGCAGCATCATGGACGTGGC
CGTTTTCCGTCTGTCGAAGCGTGACCGACGAGCTGGCGAGGTGATCCGCTACGAGCTTCCAGACGGGC
ACGTAGAGGTTTCCGCAGGGCCGGCCGGCATGGCCAGTGTGTGGGATTACGACCTGGTACTGATGGCG
GTTTCCCATCTAACCGAATCCATGAACCGATACCGGGAAGGGAAGGGAGACAAGCCCGGCCGCGTGTT

CCGTCCACACGTTGCGGACGTACTCAAGTTCTGCCGGCGAGCCGATGGCGGAAAGCAGAAAGACGACC
TGGTAGAAACCTGCATTCGGTTAAACACCACGCACGTTGCCATGCAGCGTACGAAGAAGGCCAAGAAC
GGCCGCCTGGTGACGGTATCCGAGGGTGAAGCCTTGATTAGCCGCTACAAGATCGTAAAGAGCGAAAC
CGGGCGGCCGGAGTACATCGAGATCGAGCTAGCTGATTGGATGTACCGCGAGATCACAGAAGGCAAGA
ACCCGGACGTGCTGACGGTTCACCCCGATTACTTTTTGATCGATCCCGGCATCGGCCGTTTTCTCTAC
CGCCTGGCACGCCGCGCCGCAGGCAAGGCAGAAGCCAGATGGTTGTTCAAGACGATCTACGAACGCAG
TGGCAGCGCCGGAGAGTTCAAGAAGTTCTGTTTCACCGTGCGCAAGCTGATCGGGTCAAATGACCTGC
CGGAGTACGATTTGAAGGAGGAGGCGGGGCAGGCTGGCCCGATCCTAGTCATGCGCTACCGCAACCTG
ATCGAGGGCGAAGCATCCGCCGGTTCCTAATGTACGGAGCAGATGCTAGGGCAAATTGCCCTAGCAGG
GGAAAAAGGTCGAAAAAGCTTCTTTCCTGTGGATAGCACGTACATTGGGAACCCAAAGCCGTACATTG
GGAACCGGAACCCGTACATTGGGAACCCAAAGCCGTACATTGGGAACCGGTCACACATGTAAGTGACT
GATATAAAAGAGAAAAAAGGCGATTTTTCCGCCTAAAACTCTTTAAAACTTATTAAAACTCTTAAAAC
CCGCCTGGCCTGTGCATAACTGTCTGGCCAGCGCACAGCCGAACAGCTGCAAAAAGCGCCTACCCTTC
GGTCGCTGCGCTCCCTACGCCCCGCCGCTTCGCGTCGGCCTATCGCGGCCGCTGGCCGCTCAAAAATG
GCTGGCCTACGGCCAGGCAATCTACCAGGGCGCGGACAAGCCGCGCCGTCGCCACTCGACCGCCGGCG
CCCACATCAAGGCTCCGAGTGCGCGGAACCCCTATTTGTTTATTTTTCTAAATACATTCAAATATGTA
TCCGCTCATGAGACAATAACCCTGATAAATGCTTCAATAATATTGAAAAAGGAAGAGTATGGCTAAAA
TGAGAATATCACCGGAATTGAAAAAACTGATCGAAAAATACCGCTGCGTAAAAGATACGGAAGGAATG
TCTCCTGCTAAGGTATATAAGCTGGTGGGAGAAAATGAAAACCTATATTTAAAAATGACGGACAGCCG
GTATAAAGGGACCACCTATGATGTGGAACGGGAAAAGGACATGATGCTATGGCTGGAAGGAAAGCTGC
CTGTTCCAAAGGTCCTGCACTTTGAACGGCATGATGGCTGGAGCAATCTGCTCATGAGTGAGGCCGAT
GGCGTCCTTTGCTCGGAAGAGTATGAAGATGAACAAAGCCCTGAAAAGATTATCGAGCTGTATGCGGA
GTGCATCAGGCTCTTTCACTCCATCGACATATCGGATTGTCCCTATACGAATAGCTTAGACAGCCGCT
TAGCCGAATTGGATTACTTACTGAATAACGATCTGGCCGATGTGGATTGCGAAAACTGGGAAGAGGAC
ACTCCATTTAAAGATCCGCGCGAGCTGTATGATTTTTTAAAGACGGAAAAGCCCGAAGAGGAACTTGT
CTTTTCCCACGGCGACCTGGGAGACAGCAACATCTTTGTGAAAGATGGCAAAGTAAGTGGCTTTATTG
ATCTTGGGAGAAGCGGCAGGGCGGACAAGTGGTATGACATTGCCTTCTGCGTCCGGTCGCTCAGGGAG
GATATCGGGGAAGAACAGTATGTCGAGCTATTTTTTGACTTACTGGGGATCAAGCCTGATTGGGAGAA
AATAAAATATTATATTTTACTGGATGAATTGTTTTAGCTGTCAGACCAAGTTTACTCATATATACTTT
AGATTGATTTAAAACTTCATTTTTAATTTAAAAGGATCTAGGTGAAGATCCTTTTTGATAATCTCATG
ACCAAAATCCCTTAACGTGAGTTTTCGTTCCACTGAGCGTCAGACCCCGTAGAAAAGATCAAAGGATC
TTCTTGAGATCCTTTTTTTCTGCGCGTAATCTGCTGCTTGCAAACAAAAAAACCACCGCTACCAGCGG
TGGTTTGTTTGCCGGATCAAGAGCTACCAACTCTTTTTCCGAAGGTAACTGGCTTCAGCAGAGCGCAG
ATACCAAATACTGTTCTTCTAGTGTAGCCGTAGTTAGGCCACCACTTCAAGAACTCTGTAGCACCGCC
TACATACCTCGCTCTGCTAATCCTGTTACCAGTGGCTGCTGCCAGTGGCGATAAGTCGTGTCTTACCG
GGTTGGACTCAAGACGATAGTTACCGGATAAGGCGCAGCGGTCGGGCTGAACGGGGGGTTCGTGCACA
CAGCCCAGCTTGGAGCGAACGACCTACACCGAACTGAGATACCTACAGCGTGAGCTATGAGAAAGCGC
CACGCTTCCCGAAGGGAGAAAGGCGGACAGGTATCCGGTAAGCGGCAGGGTCGGAACAGGAGAGCGCA
CGAGGGAGCTTCCAGGGGGAAACGCCTGGTATCTTTATAGTCCTGTCGGGTTTCGCCACCTCTGACTT
GAGCGTCGATTTTTGTGATGCTCGTCAGGGGGGCGGAGCCTATGGAAAAACGCCAGCAACGCGGCCTT
TTTACGGTTCCTGCTCGGATCTGTTGGACCGGACAGTAGTCATGGTTGATGGGCTGCCTGTATCGAGT
GGTGATTTTGTGCCGAGCTGCCGGTCGGGGAGCTGTTGGCTGG

>2_TYPEX_003
GAACACTCTGtgccgaattcggatccagcgtcgatctagtaacatagatgacaccgcgcgcgataatt
tatcctagtttgcgcgctatattttgttttctatcgcgtattaaatgtataattgcgggactctaatc
ataaaaacccatctcataaataacgtcatgcattacatgttaattattacatgcttaacgtaattcaa
cagaaattatatgataatcatcgcaagaccggcaacaggattcaatcttaagaaactttattgccaaa
tgtttgaacgatctgcttgacaagcctattcctttgccctcggacgagtgctggggcgtcggtttcca
ctatcggcgagtacttctacacagccatcggtccagacggccgcgcttctgcgggcgatttgtgtacg
cccgacagtcccggctccggatcggacgattgcgtcgcatcgaccctgcgcccaagctgcatcatcga

aattgccgtcaaccaagctctgatagagttggtcaagaccaatgcggagcatatacgcccggagccgc
ggcgatcctgcaagctccggatgcctccgctcgaagtagcgcgtctgctgctccatacaagccaacca
cggcctccagaagaagatgttggcgacctcgtattgggaatccccgaacatcgcctcgctccagtcaa
tgaccgctgttatgcggccattgtccgtcaggacattgttggagccgaaatccgcgtgcacgagatgc
cggacttcggggcagtcctcggcccaaagcatcagctcatcgagagcctgcgcgacggacgcactgac
ggtgtcgtccatcacagtttgccagtgatacacatggggatcagcaatcgcgcatatgaaatcacgcc
atgtagtgtattgaccgattccttgcggtccgaatgggccgaacccgctcgtctggctaagatcggcc
gcagcgatcgcatccatggcctccgcgaccggctgcagttatcatcatcatagacacacgaaata
aagtaatcagattatcagttaaagctatgtaatatttacaccataaccaatcaattaaaaaatagatc
agtttaaagaaagatcaaagctcaaaaaaataaaaagagaaaagggtcctaaccaagaaaatgaagga
gaaaaactagaaatttacctgcagaacagcgggcagttcggtttcaggcaggtcttgcaacgtgacac
cctgggcacggcgggagatgcaataggtcaggctctcgctgaattccccaatgtcaagcacttccgga
atcgggagcgcggccgatgcaaagtgccgataaacataacgatctttgtagaaaccatcggcgcagct
atttacccgcaggacatatccacgccctcctacatcgaagctgaaagcacgagattcttcgccctccg
agagctgcatcaggtcggacacgctgtcgaacttttcgatcagaaacttctcgacagacgtcgcggtg
agttcaggctttttcattgcgtgtcctctccaaatgaaatgaacttccttatatagaggaagggtctt
gcgaaggatagtgggattgtgcgtcatcccttacgtcagtggagatgtcacatcaatccacttgcttt
gtagacgtggttggaacctcttctttttccacgatgctcctcgtgggtgggggtccatctttgggacc
actgtcggcagagagatcttgaatgatagcctttcctttatcgcaatgatggcatttgtaggagccac
cttccttttctactgtcctttcgatgaagtgacagatagctgggcaatggaatccgaggaggtttccc
gaaattatcctttgttgaaaagtctcaatagccctttgatcttctgagactgtatctttgacatttttt
ggagtagaccagagtgtcgtgctccaccatgttgacctccGCAAGAATTCAAGCTTAGCGatctggat
tttagtactggattttggttttaggaattagaaattttattgatagaagtattttacaaatacaaata
catactaagggtttcttatatgctcaacacatgagcgaaaccctataggaaccctaattcccttatct
gggaactactcacacattattatggagaaactcgagcttgtcgatcgactctagctagagaagcCTAG
AGCGGTTGATCTGTGAGGTCTGGCCTTGTGCCATGCGCTTCCGCGAGAATATCCAGACAGTTGTGTTG
GAGGCCCTCCTCTGGGAGTGGCAGCAGTGTCGCCGGATTGAGGGCGACCACTGGGCCGAACTGGACGC
GATCGGTGTCGAGGAGGAGCGCTTGGTAATGGGTCATGCGCGCATTGGAGAGCCACCTATCCGGCGGC
TGTTTGACGAGGGCTTCCACGGCATGTGGCGCGAGGATCACCAGCGGCTGGCCCATTGTGAGTTTGCC
GGCATCCTTTGTGAGCACGGCAATCGCGGCGACCATGCGCAGACACGGTGGCCAGCCGGCGGCCACTG
GATCGAGTTTTTTGGAGAGATAGGCCACTGGCCTCCTCCACGGGCCGAGCTTCTGTGTGAGCACGCCC
TTCGCGTAGCCTTGCTTCTCATCGACGAACAGCTCGAACGGCTTTGTGAGATCTGGGAGCCCGAGCGC
TGGGGCGGTGAGGAGCGCTTGTTTGATCTCTTGGTAGGCTTTCTGCTGATCCGGCCCCCAGTTAAACA
GTGTGCCCGGCTTGGTGAGTGGGTAGAGTGGCGCGGCCATCTCGGCAAAGCCCGGGATGAAGAGGCGA
CAGAAGCCGGCTTTGCCGAGAAACTCGCGCAGCTGGCGTGGTGTTTTCGGTGTCGGTTGCCCCATCAC
TGTTTCTTTGCGCGCTTCGGTGAGCCAGCGTTGGCCTTCCTTGAGCAGATACCCGAGGTACTTCACTT
GCTTCTGGCAGATTTGGGCCTTTTTCGCGGACGCGCGATAGCCGAGGTTCCCGAGTGTTTGGAGGAGC
GCGCGTGTGCCTTGTTGACAATCCAGCTCGGAGGTCGCGGCGAGGAGGAGATCATCCACATACTGGAG
GAGGATGAGGTCCGGATGCTGGATGCGAAAATCCGCGAGGTCCCTATGGAGCGCCTCGTTGAAGAGTG
TCGGGCTGTTCTTGAAGCCTTGCGGGAGGCGTGTCCATGTCAGTTGGCCGGAGATGCCCATCTCCGGG
TCGCGCCACTCGAAGGCGAAGAGCGGCTGGCTTGTTGGGTGGAGGCGGAGACAGAAGAAGGCGTCTTT
GAGGTCCAGCACTGTATACCACTGGTGGGATGGTGGGAGCCCGGAGAGCAGATTATACGGGTTCGGGA
CGGTCGGGTGGATaTCTTCCACGCGCTTGTTGACCTCGCGGAGATCTTGGACCGGGCGATAGTCGTTG
GTGCCCGGCTTTTTGACCGGGAGGAGTGGTGTATTCCACGGGGATTGGCACGGCACGAGAATGCCTTG
ATCGAGGAGGCGCTGAATGTGCGGCTTAATCCCGAGCCTCGCCTCTTGGCTCATCGGGTACTGCTTGA
TGGACACCGGTGTGGAGGTCGCCTTCAGCGGGATAATGAGTGGGGCTTGGCGGACGGCGAGGCCCATG
CCGCCGGTTTCGGCCCACGCTTGTGGGAAGTCGCTGAGCCATGTGCTGCCGAGGGACACGTCCGGTTC
TTTGGAGGTTTCGTGGAGGCGATACTCATCCTCGATGTTGAGGGTAGGCCTAGATCCTCCGGAAGAGC
CTCCGCTAGATTCTGGTGTAGCGCTTTCGGATGTTCCTGGTGTCTCGGAGCCGCTAGATCCTCCGGAA
GATCCTCCGCTGAGCTCCACCTTCCGCTTCTTCTTTGGCGAACCAAGGAGGGATGTTTGTGGCCTTGG
GCCGCGTGGGCCGCGCGGCTTTTTCGGGCAGTCTTTGGCCCAGTGGCCCTTCTCCTTGCAGTAGGCAC
ACTGATCCCTATCGAGCTGGGACCTTCTGCGTTCTCCGCCCTGGCGGTCCTGCTTTTGGCCGGACACC
ACTGTGGCAGATTCTGGGGTGGCCGACTCGGAGGTGCCTGGCGTTTCGCTCCCGGAACCTGAATCACC
ACCGAGCTGTGAGAGATCGATCCTAGTCTCGTAGAGTCCAGTGATAGACTGATGGATGAGGGTAGCAT
CGAGCACTTCTTTGGTAGAGGTGTATCTCTTCCTATCGATGGTTGTATCGAAGTACTTGAAAGCAGCA
GGAGCACCGAGGTTGGTAAGGGTGAAGAGATGGATGATGTTCTCTGCCTGTTCCCTGATAGGCTTATC

TCTGTGCTTGTTGTAAGCAGACAACACCTTATCGAGGTTTGCATCAGCGAGGATCACCCTTTTAGAGA
ACTCAGAGATCTGCTCGATGATCTCATCCAAGTAGTGCTTGTGCTGCTCAACGAAAAGTTGCTTCTGC
TCGTTATCTTCTGGAGATCCCTTCAACTTCTCGTAGTGAGAAGCGAGGTAAAGAAAGTTAACGTACTT
AGATGGGAGAGCAAGCTCGTTTCCCTTTTGAAGCTCACCAGCAGAAGCGAGCATCCTCTTTCTACCGT
TCTCGAGTTCGAAGAGTGAGTACTTTGGGAGCTTGATGATGAGATCCTTCTTAACCTCTTTGTATCCC
TTAGCCTCGAGGAAATCGATTGGGTTCTTCTCGAAAGATGACCTTTCCATGATAGTGATTCCGAGAAG
TTCCTTAACAGACTTGAGCTTCTTACTCTTTCCCTTCTCAACCTTAGCCACAACGAGAACAGAGTAAG
CCACGGTAGGAGAATCGAAACCACCGTATTTCTTAGGGTCCCAATCCTTCTTCCTAGCAATGAGCTTA
TCAGAGTTCCTCTTAGGGAGGATAGACTCTTTAGAGAATCCACCGGTCTGCACCTCGGTTTTCTTAAC
GATGTTCACCTGTGGCATAGAGAGCACCTTTCTAACGGTAGCGAAATCCCTTCCCTTATCCCACACGA
TCTCACCTGTTTCACCGTTTGTCTCGATGAGTGGCCTCTTTCTGATCTCACCGTTAGCGAGGGTAATC
TCGGTCTTGAAGAAATTCATGATGTTAGAGTAGAAGAAATACTTAGCGGTAGCCTTTCCGATCTCTTG
CTCAGACTTAGCGATCATCTTCCTCACATCGTACACCTTGTAATCACCGTACACGAACTCTGACTCGA
GCTTAGGATACTTCTTGATGAGAGCGGTTCCAACAACAGCGTTAAGGTAAGCATCGTGAGCGTGGTGG
TAGTTGTTGATTTCCCTCACCTTGTAGAATTGGAAATCCTTTCTGAAATCAGACACGAGCTTTGACTT
GAGGGTGATAACCTTCACTTCCCTGATCAACTTATCGTTCTCATCGTACTTGGTGTTCATCCTAGAAT
CGAGGATCTGTGCAACGTGCTTAGTGATCTGCCTGGTTTCCACAAGCTGCCTCTTGATGAATCCTGCC
TTATCCAATTCAGAGAGTCCTCCCCTCTCAGCCTTAGTCAAGTTATCGAACTTTCTCTGAGTGATGAG
CTTAGCGTTGAGGAGCTGCCTCCAATAGTTCTTCATTTCTTCACAACCTCTTCACTTGGCACGTTAT
CACTCTTACCCCTGTTCTTATCAGACCTGGTGAGCACCTTGTTATCGATAGAATCATCCTTCAAGAAT
GACTGTGGCACGATAGCATCAACATCGTAATCAGAGAGCCTGTTGATATCCAACTCTTGATCCACATA
CATATCCCTTCCGTTCTGGAGGTAGTAGAGGTAGAGCTTCTCATTCTGGAGCTGAGTGTTCTCAACAG
GGTGCTCTTTGAGGATCTGAGATCCAAGCTCTTTGATACCTTCCTCGATCCTCTTCATCCTTTCCCTA
GAGTTCTTCTGTCCCTTCTGAGTGGTCTGGTTCTCTCTAGCCATTTCGATCACGATGTTCTCAGGCTT
ATGCCTTCCCATCACCTTCACCAACTCATCCACAACCTTCACAGTCTGGAGGATTCCCTTCTTGATTG
CAGGAGATCCAGCGAGGTTAGCGATATGCTCATGGAGACTATCACCCTGTCCTGAAACCTGAGCCTTC
TGGATATCCTCTTTAAAGGTGAGAGAATCATCGTGGATGAGCTGCATGAAGTTTCTGTTAGCGAATCC
ATCAGACTTGAGGAAATCAAGGATTGTCTTTCCAGACTGCTTATCCCTGATTCCGTTAATGAGCTTTC
TTGAGAGCCTTCCCCAACCAGTGTATCTTCTTCTCTTCAACTGCTTCATCACCTTATCATCGAAGAGA
TGAGCGTAGGTCTTGAGCCTTTCTTCAATCATCTCTCTATCTTCAAAGAGGGTGAGGGTAAGAACGAT
ATCCTCCAAGATATCCTCGTTTTCCTCGTTATCCAAGAAATCCTTATCCTTAATGATCTTGAGGAGAT
CGTGGTAGGTTCCGAGAGATGCGTTGAACCTATCCTCAACACCAGAAATCTCAACTGAATCGAAGCAC
TCGATTTTCTTGAAGTAATCCTCTTTGAGCTGCTTCACGGTCACCTTTCTGTTGGTCTTGAACAAGAG
ATCAACGATAGCCTTCTTTTGCTCACCTGACAAAAAAGCAGGCTTCCTCATTCCCTCGGTCACGTACT
TAACCTTGGTCAACTCGTTGTACACGGTGAAGTACTCGTAGAGCAAAGAGTGCTTAGGGAGCACCTTC
TCGTTTGGAAGGTTCTTATCGAAGTTGGTCATCCTCTCGATGAAAGACTGAGCACTAGCACCCTTATC
CACCACCTCTTCGAAGTTCCAAGGGGTGATGGTTTCCTCAGACTTTCTGGTCATCCAAGCGAATCTTG
AGTTTCCTCTAGCGAGAGGTCCCACGTAGTAAGGGATTCTGAAGGTGAGAATCTTCTCAATCTTTTCC
CTGTTATCCTTGAGGAATGGGTAGAAATCCTCTTGCCTTCTAAGGATAGCGTGCAACTCTCCGAGGTG
GATCTGATGAGGGATAGATCCGTTATCGAAGGTCCTCTGCTTTCTGAGAAGATCCTCTCTATTGAGCT
TCACGAGGAGTTCCTCGGTTCCATCCATCTTCTCGAGGATAGGCTTGATGAACTTGTAGAACTCTTCT
TGAGATGCACCACCATCGATGTAACCAGCGTATCCGTTCTTAGACTGATCGAAGAAAATCTCTTTGTA
CTTCTCTGGGAGCTGCTGTCTAACAAGAGCCTTGAGAAGTGTGAGATCCTGGTGGTGCTCATCGTATC
TCTTGATCATAGAAGCTGAGAGTGGAGCCTTGGTGATCTCGGTGTTCACTCTGAGGATATCACTGAGG
AGGATAGCATCAGAGAGGTTCTTAGCAGCGAGGAACAAATCAGCGTACTGATCTCCGATCTGAGCGAG
GAGGTTATCGAGATCATCATCGTAGGTATCCTTTGAGAGCTGGAGCTTTGCATCCTCAGCGAGATCGA
AGTTAGACTTGAAGTTAGGGGTGAGTCCGAGAGAGAGAGCGATCAAGTTTCCGAAAAGTCCGTTCTTC
TTCTCACCAGGGAGCTGAGCAATGAGGTTCTCAAGCCTTCTTGACTTAGAGAGCCTAGCAGAGAGGAT
AGCCTTAGCATCCACACCTGAAGCGTTGATAGGGTTCTCTTCGAAAAGCTGGTTGTAGGTCTGCACGA
GCTGGATGAACAACTTATCCACATCAGAGTTATCAGGGTTGAGATCACCCTCGATGAGGAAGTGTCCT
CTGAACTTGATCATGTGAGCGAGAGCGAGGTAGATGAGCCTGAGATCAGCCTTATCAGTAGAATCAAC
GAGCTTCTTTCTGAGGTGGTAGATAGTAGGGTACTTCTCGTGGTATGCCACCTCATCAACGATGTTTC
CGAAGATAGGGTGCCTCTCGTGCTTCTTATCTTCTTCCACGAGGAATGACTCTTCGAGCCTGTGGAAG
AATGAATCATCCACTTTAGCCATCTCGTTAGAGAAGATCTCTTGGAGGTAGCAGATCCTGTTCTTTCT
TCTGGTGTACCTTCTTCTAGCGGTTCTCTTGAGTCTGGTAGCCTCAGCAGTTTCACCAGAATCGAAGA
GGAGAGCACCGATAAGGTTTTTCTTGATAGAGTGCCTATCGGTGTTTCCGAGAACCTTGAACTTCTTA

GATGGCACCTTGTACTCATCGGTGATCACAGCCCATCCCACAGAGTTAGTTCCGATATCGAGTCCGAT
AGAGTACTTCTTATCAACCTTTCTCTTCTTCTTAGGAGATTCAAATTCAGATCCATCAGCAGTTCTCT
TCATCATTcaacctttctgcaggcacatcaatactgttgaggcaacacccatagaagcacggaaccaa
ggagcaccgacgtccatagcacacagccaacctcgctgaccttcgactcctcttcaagtcaaggatag
acaaatctatcatgcaggaaacttacgaggcacctccgcaccagtgaccgaactgaacaccttaccga
gataacaggactgaccaaagtgggatgtacttggctaaaacgaacctaaaaaatcactcgctcgaggc
cccaatcgacaccacgtaaaagctccagtcgggtttcaggtctagacaatcacaatatcgagcaatcg
taacgacgtacagtgacgagggcacaaggcgcataccagtagacaagatgcagctcttcgatgtcgat
gctctccacgcaagagagaaagaagtgaaaggaactctagggttgcggcgatcgattttaagccagac
gctccactgcctcccagcgagctttgagagcaagcgcgaccacagcactagacgccaaagtagaagaa
attagggaatgattagcgaaattgcggagggttttgatataattaaatacctaggtctgccccgttta
taataacgagcgttcgataaggactttgcatcggctccgttcaagttattgcgacttagggtcggcgg
ccggccattcgcagataggcggccaattcagcatgcgagggcaagttgggaggccgacgacgaagggt
taggaatattcaaagcctcttttccatttatatattgccctgtccatgccccttcggactctttgtcg
tgactgccggccagcatcgtcatccgaattgctcacactcctccgtctgtcacatcgtccgtgcttag
ggccggcacgccctttcaatttaacattccaaaataagatgacgtaggtttagcgaacccgactgctg
ataccacaccgattgctccggtttttcgagtaccttttccgcgcttattacaagctgcgagcaaactt
tctagaaagtagagaggtggccgaagtcgcttttggcctcccgcgcctttctccttctgcacatactt
ccaaacggtggactagcatcccatagatccatgacatgcacgcaatcacatgagatttactgttgct
ttcttcttctggcctcaccgtcctccccgtcctcccttcgctttttgggacgcgggaaaacctagaag
gcttctgattgaagtttccttttctgcttgtcggcctttacactttggagaacgaagttctaatctct
cttgaagttgaccaaatattagtatagtgaactgatcgagaagatgcgggtccgatccaataggagag
aacaaaatattacgagacagaaagagataattaagcaatttatatgcactttaaaatggaatcttttc
gattatcatcaatgagcattatattgtgaaacggatttaaaggattagctattttgaaaggtttttt
aatctgtttttaagatctttagcaaattctttttaagttacccaaatatcgatacccgttaacaatc
aaaaatcgatgacagagttgtgaatcatcatcaagatctgcaaatttgatatgattttgaaaatttga
gatacaaatattgttaaaataatgcatttaactgggaaatctatcaacttaccttacaattgagttat
tttgatatgtctctcaacaatgtgtgtgactcatttgctccactagaattcgagctcagcgaaaaaaa
gcaccgactcggtgccactttttcaagttgataacggactagccttattttaacttgctatttctagc
tctaaaacagacataaaaaacaaaaaaaTTTCTAGTTGGTTTAACGCGTAACTAGATAGAACCGCGTC
AAGAGAGAGAGGTACCAATCATCCTTCTTAGTACTTGACGCACCGACTCGGTGCCACTTTTTCAAGTT
GATAACGGACTAGCCTTATTTTAACTTGCTATTTCTAGCTCTAAAACAGAAGGATGATTGGTACCGAc
TATGgcaaggtgcaatttacggatttatacactacatttgcattgtgcgttttgcttatagttttact
ctcgataagcaaaccacgtggtcccttctccgtcattgctgtcgtgaatttctgaagtaaccggaaaa
tgtttcttggccgtcatgagttcaatcctgctgtgcgatcgcgttctggctcggtccaataccactcc
gtctgctcgtttcggtatatatatcggacctagaggagaatggagatcgtcaagacatcccaccatgt
ccacaaatttgtgctccatcgcgggtttctttccatcgtgaaagcgatgcatacaccaaaaaaacgaag
cataactgcgcatgccccaagagcatcgtttagagcaacaggaataagacttagatttgattcgatat
cgtaaagatagcgaccatgggacacaccacttgtttctagaacttcgaaccccgggggaaaatagttg
atggggccttcccatcctttttgctatattacttggtattcccgaagaagggcttctttggacggaga
ggagtgtatgtacgatgctagctccgtgttttcgtgttttcttgtttccccaaacctgaaggatttgg
gtggaagtaccaaggagattcaactgtccagttgtgcgggctacccaaatgaaagaactcccacacgc
gcaaatttgtttcctacattgagaaataactgcaaaagcctgcacgttgctgtaagaaaaataatctg
taggaaattaggaaactacagagctcgtctgaaaattcccagctttcatttcaggtttacggtatttt
atttctgacgtgacagaaataaaattggaaattggaaattcattggatgcatggccaggtatcggtgc
agtcacatctttcgcacccgttttatcactacaacgttagtgcggtgtacatttcaatcgctgttgtt
atctgagcttatctcgccgttccatttgagtcatccgatccaacgagaacagacacgtctgacgcagt
cctttagaatccaggtcatttcgcattcttccccacaggaatcaagtctctaatcttgttccgatccg
tcagacaacacttgctccgtccgcagagccctgtgcgattgagtgccgagtgttcttggtgcttcatt
ttagtaaaactcgatcagcgatcgttcttggttctggacagatctgcaggtcacgagaacagctcatg
tcttttatccttagagaaatcggtcatatgtttgcagcttgtcagaacttttgcgactttattgatta
cgtgacatacgcggaacaagaggctacacaatcggaagggcgaatcggccggatgcactggcagtgga
ttccatcaagacaatacagtcgcttgcggaaacagcattcacatttgttgtcaatgcagctaggaaga
ggtaaattcacatcttgacatgctaaacaaatcagcacctttgaacataattccgttttgtatgtcaa
tacaaaagtgtactactaagccattgctaaaatgcataaatagcaatgctttcctctgagaacaaact
gattgccactaatttaatgtgaacatggattacaatagagcctgtagctcaaatacctgcgcatccct

acatcatgcagcaaaattggatagcatccgctagaacgagatttagggcgctggagctggactgatgg
gtcctttctacttcctgatggtggggacggaggaattacaacccaaggtgtccggccggtaggagcc
tttctgggtggtgggggagccgtggccagttcagtcacacgaagctgccggctggtccctactgccac
ctctgtcatgtccacgggaacatcgactgaagctttctcttgctttatcacgtccactttttcgtga
gtttacgtcccggaactaatgctctctctgtgaaagtcgtccccttgacctgccatacatacatctcg
actgaacaacttcagccatcactccTTACGAGGATGCACATGTGACCGAGGGACACGAAGTGATCCGT
TTAAACTATCAGTGTTTGACAGGATATATTGGCGGGTAAACCTAAGAGAAAAGAGCGTTTATTAGAAT
AATCGGATATTTAAAAGGGCGTGAAAAGGTTTATCCGTTCGTCCATTTGTATGTGCATGCCAACCACA
GGGTTCCCCTCGGGAGTCAGCCGTGCGGCTGCATGAAATCCTGGCCGGTTTGTCTGATGCCAAGCTGG
CGGCCTGGCCGGCCAGCTTGGCCGCTGAAGAAACCGAGCGCCGCCGTCTAAAAAGGTGATGTGTATTT
GAGTAAAACAGCTTGCGTCATGCGGTCGCTGCGTATATGATGCGATGAGTAAATAAACAAATACGCAA
GGGGAACGCATGAAGGTTATCGCTGTACTTAACCAGAAAGGCGGGTCAGGCAAGACGACCATCGCAAC
CCATCTAGCCCGCGCCCTGCAACTCGCCGGGGCCGATGTTCTGTTAGTCGATTCCGATCCCCAGGGCA
GTGCCCGCGATTGGGCGGCCGTGCGGGAAGATCAACCGCTAACCGTTGTCGGCATCGACCGCCCGACG
ATTGACCGCGACGTGAAGGCCATCGGCCGGCGCGACTTCGTAGTGATCGACGGAGCGCCCCAGGCGGC
GGACTTGGCTGTGTCCGCGATCAAGGCAGCCGACTTCGTGCTGATTCCGGTGCAGCCAAGCCCTTACG
ACATATGGGCCACCGCCGACCTGGTGGAGCTGGTTAAGCAGCGCATTGAGGTCACGGATGGAAGGCTA
CAAGCGGCCTTTGTCGTGTCGCGGGCGATCAAAGGCACGCGCATCGGCGGTGAGGTTGCCGAGGCGCT
GGCCGGGTACGAGCTGCCCATTCTTGAGTCCCGTATCACGCAGCGCGTGAGCTACCCAGGCACTGCCG
CCGCCGGCACAACCGTTCTTGAATCAGAACCCGAGGGCGACGCTGCCCGCGAGGTCCAGGCGCTGGCC
GCTGAAATTAAATCAAAACTCATTTGAGTTAATGAGGTAAAGAGAAAATGAGCAAAAGCACAAACACG
CTAAGTGCCGGCCGTCCGAGCGCACGCAGCAGCAAGGCTGCAACGTTGGCCAGCCTGGCAGACACGCC
AGCCATGAAGCGGGTCAACTTTCAGTTGCCGGCGGAGGATCACACCAAGCTGAAGATGTACGCGGTAC
GCCAAGGCAAGACCATTACCGAGCTGCTATCTGAATACATCGCGCAGCTACCAGAGTAAATGAGCAAA
TGAATAAATGAGTAGATGAATTTTAGCGGCTAAAGGAGGCGGCATGGAAAATCAAGAACAACCAGGCA
CCGACGCCGTGGAATGCCCCATGTGTGGAGGAACGGGCGGTTGGCCAGGCGTAAGCGGCTGGGTTGTC
TGCCGGCCCTGCAATGGCACTGGAACCCCCAAGCCCGAGGAATCGGCGTGACGGTCGCAAACCATCCG
GCCCGGTACAAATCGGCGCGGCGCTGGGTGATGACCTGGTGGAGAAGTTGAAGGCCGCGCAGGCCGCC
CAGCGGCAACGCATCGAGGCAGAAGCACGCCCCGGTGAATCGTGGCAAGCGGCCGCTGATCGAATCCG
CAAAGAATCCCGGCAACCGCCGGCAGCCGGTGCGCCGTCGATTAGGAAGCCGCCCAAGGGCGACGAGC
AACCAGATTTTTTCGTTCCGATGCTCTATGACGTGGGCACCCGCGATAGTCGCAGCATCATGGACGTG
GCCGTTTTCCGTCTGTCGAAGCGTGACCGACGAGCTGGCGAGGTGATCCGCTACGAGCTTCCAGACGG
GCACGTAGAGGTTTCCGCAGGGCCGGCCGGCATGGCCAGTGTGTGGGATTACGACCTGGTACTGATGG
CGGTTTCCCATCTAACCGAATCCATGAACCGATACCGGGAAGGGAAGGGAGACAAGCCCGGCCGCGTG
TTCCGTCCACACGTTGCGGACGTACTCAAGTTCTGCCGGCGAGCCGATGGCGGAAAGCAGAAAGACGA
CCTGGTAGAAACCTGCATTCGGTTAAACACCACGCACGTTGCCATGCAGCGTACGAAGAAGGCCAAGA
ACGGCCGCCTGGTGACGGTATCCGAGGGTGAAGCCTTGATTAGCCGCTACAAGATCGTAAAGAGCGAA
ACCGGGCGGCCGGAGTACATCGAGATCGAGCTAGCTGATTGGATGTACCGCGAGATCACAGAAGGCAA
GAACCCGGACGTGCTGACGGTTCACCCCGATTACTTTTTGATCGATCCCGGCATCGGCCGTTTTCTCT
ACCGCCTGGCACGCCGCGCCGCAGGCAAGGCAGAAGCCAGATGGTTGTTCAAGACGATCTACGAACGC
AGTGGCAGCGCCGGAGAGTTCAAGAAGTTCTGTTTCACCGTGCGCAAGCTGATCGGGTCAAATGACCT
GCCGGAGTACGATTTGAAGGAGGAGGCGGGGCAGGCTGGCCCGATCCTAGTCATGCGCTACCGCAACC
TGATCGAGGGCGAAGCATCCGCCGGTTCCTAATGTACGGAGCAGATGCTAGGGCAAATTGCCCTAGCA
GGGGAAAAAGGTCGAAAAAGCTTCTTTCCTGTGGATAGCACGTACATTGGGAACCCAAAGCCGTACAT
TGGGAACCGGAACCCGTACATTGGGAACCCAAAGCCGTACATTGGGAACCGGTCACACATGTAAGTGA
CTGATATAAAAGAGAAAAAAGGCGATTTTTCCGCCTAAAACTCTTTAAAACTTATTAAAACTCTTAAA
ACCCGCCTGGCCTGTGCATAACTGTCTGGCCAGCGCACAGCCGAACAGCTGCAAAAAGCGCCTACCCT
TCGGTCGCTGCGCTCCCTACGCCCCGCCGCTTCGCGTCGGCCTATCGCGGCCGCTGGCCGCTCAAAAA
TGGCTGGCCTACGGCCAGGCAATCTACCAGGGCGCGGACAAGCCGCGCCGTCGCCACTCGACCGCCGG
CGCCCACATCAAGGCTCCGAGTGCGCGGAACCCCTATTTGTTTATTTTTCTAAATACATTCAAATATG
TATCCGCTCATGAGACAATAACCCTGATAAATGCTTCAATAATATTGAAAAAGGAAGAGTATGGCTAA
AATGAGAATATCACCGGAATTGAAAAAACTGATCGAAAAATACCGCTGCGTAAAAGATACGGAAGGAA
TGTCTCCTGCTAAGGTATATAAGCTGGTGGGAGAAAATGAAAACCTATATTTAAAAATGACGGACAGC
CGGTATAAAGGGACCACCTATGATGTGGAACGGGAAAAGGACATGATGCTATGGCTGGAAGGAAAGCT
GCCTGTTCCAAAGGTCCTGCACTTTGAACGGCATGATGGCTGGAGCAATCTGCTCATGAGTGAGGCCG
ATGGCGTCCTTTGCTCGGAAGAGTATGAAGATGAACAAAGCCCTGAAAAGATTATCGAGCTGTATGCG

GAGTGCATCAGGCTCTTTCACTCCATCGACATATCGGATTGTCCCTATACGAATAGCTTAGACAGCCG
CTTAGCCGAATTGGATTACTTACTGAATAACGATCTGGCCGATGTGGATTGCGAAAACTGGGAAGAGG
ACACTCCATTTAAAGATCCGCGCGAGCTGTATGATTTTTTAAAGACGGAAAAGCCCGAAGAGGAACTT
GTCTTTTCCCACGGCGACCTGGGAGACAGCAACATCTTTGTGAAAGATGGCAAAGTAAGTGGCTTTAT
TGATCTTGGGAGAAGCGGCAGGGCGGACAAGTGGTATGACATTGCCTTCTGCGTCCGGTCGCTCAGGG
AGGATATCGGGGAAGAACAGTATGTCGAGCTATTTTTTGACTTACTGGGGATCAAGCCTGATTGGGAG
AAAATAAAATATTATATTTTACTGGATGAATTGTTTTAGCTGTCAGACCAAGTTTACTCATATATACT
TTAGATTGATTTAAAACTTCATTTTTAATTTAAAAGGATCTAGGTGAAGATCCTTTTTGATAATCTCA
TGACCAAAATCCCTTAACGTGAGTTTTCGTTCCACTGAGCGTCAGACCCCGTAGAAAAGATCAAAGGA
TCTTCTTGAGATCCTTTTTTTCTGCGCGTAATCTGCTGCTTGCAAACAAAAAAACCACCGCTACCAGC
GGTGGTTTGTTTGCCGGATCAAGAGCTACCAACTCTTTTTCCGAAGGTAACTGGCTTCAGCAGAGCGC
AGATACCAAATACTGTTCTTCTAGTGTAGCCGTAGTTAGGCCACCACTTCAAGAACTCTGTAGCACCG
CCTACATACCTCGCTCTGCTAATCCTGTTACCAGTGGCTGCTGCCAGTGGCGATAAGTCGTGTCTTAC
CGGGTTGGACTCAAGACGATAGTTACCGGATAAGGCGCAGCGGTCGGGCTGAACGGGGGGTTCGTGCA
CACAGCCCAGCTTGGAGCGAACGACCTACACCGAACTGAGATACCTACAGCGTGAGCTATGAGAAAGC
GCCACGCTTCCCGAAGGGAGAAAGGCGGACAGGTATCCGGTAAGCGGCAGGGTCGGAACAGGAGAGCG
CACGAGGGAGCTTCCAGGGGGAAACGCCTGGTATCTTTATAGTCCTGTCGGGTTTCGCCACCTCTGAC
TTGAGCGTCGATTTTTGTGATGCTCGTCAGGGGGGCGGAGCCTATGGAAAAACGCCAGCAACGCGGCC
TTTTTACGGTTCCTGCTCGGATCTGTTGGACCGGACAGTAGTCATGGTTGATGGGCTGCCTGTATCGA
GTGGTGATTTTGTGCCGAGCTGCCGGTCGGGGAGCTGTTGGCTGGCTGGTGGCAGGATATATTGTGGT
GTAAACAAATTGACGCTTAGACAACTTAATAACACATTGCGGACGTTTTTAATGTACTGGGGTT

>2_TYPEX_005
gaccgctgttatgcggccattgtccgtcaggacattgttggagccgaaatccgcgtgcacgagatgcc
ggacttcggggcagtcctcggcccaaagcatcagctcatcgagagcctgcgcgacggacgcactgacg
gtgtcgtccatcacagtttgccagtgatacacatggggatcagcaatcgcgcatatgaaatcacgcca
tgtagtgtattgaccgattccttgcggtccgaatgggccgaacccgctcgtctggctaagatcggccg
cagcgatcgcatccatggcctccgcgaccggctgcagttatcatcatcatcatagacacacgaaataa
agtaatcagattatcagttaaagctatgtaatatttacaccataaccaatcaattaaaaaatagatca
gtttaaagaaagatcaaagctcaaaaaaataaaaagagaaaagggtcctaaccaagaaaatgaaggag
aaaaactagaaatttacctgcagaacagcgggcagttcggtttcaggcaggtcttgcaacgtgacacc
ctgggcacggcgggagatgcaataggtcaggctctcgctgaattccccaatgtcaagcacttccggaa
tcgggagcgcggccgatgcaaagtgccgataaacataacgatctttgtagaaaccatcggcgcagcta
tttacccgcaggacatatccacgccctcctacatcgaagctgaaagcacgagattcttcgccctccga
gagctgcatcaggtcggacacgctgtcgaacttttcgatcagaaacttctcgacagacgtcgcggtga
gttcaggctttttcattgcgtgtcctctccaaatgaaatgaacttccttatatagaggaagggtcttg
cgaaggatagtgggattgtgcgtcatcccttacgtcagtggagatgtcacatcaatccacttgctttg
tagacgtggttggaacctcttctttttccacgatgctcctcgtgggtgggggtccatctttgggacca
ctgtcggcagagagatcttgaatgatagcctttcctttatcgcaatgatggcatttgtaggagccacc
ttccttttctactgtcctttcgatgaagtgacagatagctgggcaatggaatccgaggaggtttcccg
aaattatcctttgttgaaaagtctcaatagccctttgatcttctgagactgtatctttgacattttttg
gagtagaccagagtgtcgtgctccaccatgttgacctccGCAAGAATTCAAGCTTAGCGatctggatt
ttagtactggattttggttttaggaattagaaattttattgatagaagtattttacaaatacaaatac
atactaagggtttcttatatgctcaacacatgagcgaaaccctataggaaccctaattcccttatctg
ggaactactcacacattattatggagaaactcgagcttgtcgatcgactctagctagagaagcCTAGA
GCGGTTGATCTGTGAGGTCTGGCCTTGTGCCATGCGCTTCCGCGAGAATATCCAGACAGTTGTGTTGG
AGGCCCTCCTCTGGGAGTGGCAGCAGTGTCGCCGGATTGAGGGCGACCACTGGGCCGAACTGGACGCG
ATCGGTGTCGAGGAGGAGCGCTTGGTAATGGGTCATGCGCGCATTGGAGAGCCACCTATCCGGCGGCT
GTTTGACGAGGGCTTCCACGGCATGTGGCGCGAGGATCACCAGCGGCTGGCCCATTGTGAGTTTGCCG
GCATCCTTTGTGAGCACGGCAATCGCGGCGACCATGCGCAGACACGGTGGCCAGCCGGCGGCCACTGG
ATCGAGTTTTTTGGAGAGATAGGCCACTGGCCTCCTCCACGGGCCGAGCTTCTGTGTGAGCACGCCCT

TCGCGTAGCCTTGCTTCTCATCGACGAACAGCTCGAACGGCTTTGTGAGATCTGGGAGCCCGAGCGCT
GGGGCGGTGAGGAGCGCTTGTTTGATCTCTTGGTAGGCTTTCTGCTGATCCGGCCCCCAGTTAAACAG
TGTGCCCGGCTTGGTGAGTGGGTAGAGTGGCGCGGCCATCTCGGCAAAGCCCGGGATGAAGAGGCGAC
AGAAGCCGGCTTTGCCGAGAAACTCGCGCAGCTGGCGTGGTGTTTTCGGTGTCGGTTGCCCCATCACT
GTTTCTTTGCGCGCTTCGGTGAGCCAGCGTTGGCCTTCCTTGAGCAGATACCCGAGGTACTTCACTTG
CTTCTGGCAGATTTGGGCCTTTTTCGCGGACGCGCGATAGCCGAGGTTCCCGAGTGTTTGGAGGAGCG
CGCGTGTGCCTTGTTGACAATCCAGCTCGGAGGTCGCGGCGAGGAGGAGATCATCCACATACTGGAGG
AGGATGAGGTCCGGATGCTGGATGCGAAAATCCGCGAGGTCCCTATGGAGCGCCTCGTTGAAGAGTGT
CGGGCTGTTCTTGAAGCCTTGCGGGAGGCGTGTCCATGTCAGTTGGCCGGAGATGCCCATCTCCGGGT
CGCGCCACTCGAAGGCGAAGAGCGGCTGGCTTGTTGGGTGGAGGCGGAGACAGAAGAAGGCGTCTTTG
AGGTCCAGCACTGTATACCACTGGTGGGATGGTGGGAGCCCGGAGAGCAGATTATACGGGTTCGGGAC
GGTCGGGTGGATaTCTTCCACGCGCTTGTTGACCTCGCGGAGATCTTGGACCGGGCGATAGTCGTTGG
TGCCCGGCTTTTTGACCGGGAGGAGTGGTGTATTCCACGGGGATTGGCACGGCACGAGAATGCCTTGA
TCGAGGAGGCGCTGAATGTGCGGCTTAATCCCGAGCCTCGCCTCTTGGCTCATCGGGTACTGCTTGAT
GGACACCGGTGTGGAGGTCGCCTTCAGCGGGATAATGAGTGGGGCTTGGCGGACGGCGAGGCCCATGC
CGCCGGTTTCGGCCCACGCTTGTGGGAAGTCGCTGAGCCATGTGCTGCCGAGGGACACGTCCGGTTCT
TTGGAGGTTTCGTGGAGGCGATACTCATCCTCGATGTTGAGGGTAGGCCTAGATCCTCCGGAAGAGCC
TCCGCTAGATTCTGGTGTAGCGCTTTCGGATGTTCCTGGTGTCTCGGAGCCGCTAGATCCTCCGGAAG
ATCCTCCGCTGAGCTCCACCTTCCGCTTCTTCTTTGGCGAACCAAGGAGGGATGTTTGTGGCCTTGGG
CCGCGTGGGCCGCGCGGCTTTTTCGGGCAGTCTTTGGCCCAGTGGCCCTTCTCCTTGCAGTAGGCACA
CTGATCCCTATCGAGCTGGGACCTTCTGCGTTCTCCGCCCTGGCGGTCCTGCTTTTGGCCGGACACCA
CTGTGGCAGATTCTGGGGTGGCCGACTCGGAGGTGCCTGGCGTTTCGCTCCCGGAACCTGAATCACCA
CCGAGCTGTGAGAGATCGATCCTAGTCTCGTAGAGTCCAGTGATAGACTGATGGATGAGGGTAGCATC
GAGCACTTCTTTGGTAGAGGTGTATCTCTTCCTATCGATGGTTGTATCGAAGTACTTGAAAGCAGCAG
GAGCACCGAGGTTGGTAAGGGTGAAGAGATGGATGATGTTCTCTGCCTGTTCCCTGATAGGCTTATCT
CTGTGCTTGTTGTAAGCAGACAACACCTTATCGAGGTTTGCATCAGCGAGGATCACCCTTTTAGAGAA
CTCAGAGATCTGCTCGATGATCTCATCCAAGTAGTGCTTGTGCTGCTCAACGAAAAGTTGCTTCTGCT
CGTTATCTTCTGGAGATCCCTTCAACTTCTCGTAGTGAGAAGCGAGGTAAAGAAAGTTAACGTACTTA
GATGGGAGAGCAAGCTCGTTTCCCTTTTGAAGCTCACCAGCAGAAGCGAGCATCCTCTTTCTACCGTT
CTCGAGTTCGAAGAGTGAGTACTTTGGGAGCTTGATGATGAGATCCTTCTTAACCTCTTTGTATCCCT
TAGCCTCGAGGAAATCGATTGGGTTCTTCTCGAAAGATGACCTTTCCATGATAGTGATTCCGAGAAGT
TCCTTAACAGACTTGAGCTTCTTACTCTTTCCCTTCTCAACCTTAGCCACAACGAGAACAGAGTAAGC
CACGGTAGGAGAATCGAAACCACCGTATTTCTTAGGGTCCCAATCCTTCTTCCTAGCAATGAGCTTAT
CAGAGTTCCTCTTAGGGAGGATAGACTCTTTAGAGAATCCACCGGTCTGCACCTCGGTTTTCTTAACG
ATGTTCACCTGTGGCATAGAGAGCACCTTTCTAACGGTAGCGAAATCCCTTCCCTTATCCCACACGAT
CTCACCTGTTTCACCGTTTGTCTCGATGAGTGGCCTCTTTCTGATCTCACCGTTAGCGAGGGTAATCT
CGGTCTTGAAGAAATTCATGATGTTAGAGTAGAAGAAATACTTAGCGGTAGCCTTTCCGATCTCTTGC
TCAGACTTAGCGATCATCTTCCTCACATCGTACACCTTGTAATCACCGTACACGAACTCTGACTCGAG
CTTAGGATACTTCTTGATGAGAGCGGTTCCAACAACAGCGTTAAGGTAAGCATCGTGAGCGTGGTGGT
AGTTGTTGATTTCCCTCACCTTGTAGAATTGGAAATCCTTTCTGAAATCAGACACGAGCTTTGACTTG
AGGGTGATAACCTTCACTTCCCTGATCAACTTATCGTTCTCATCGTACTTGGTGTTCATCCTAGAATC
GAGGATCTGTGCAACGTGCTTAGTGATCTGCCTGGTTTCCACAAGCTGCCTCTTGATGAATCCTGCCT
TATCCAATTCAGAGAGTCCTCCCCTCTCAGCCTTAGTCAAGTTATCGAACTTTCTCTGAGTGATGAGC
TTAGCGTTGAGGAGCTGCCTCCAATAGTTCTTCATTTTCTTCACAACCTCTTCACTTGGCACGTTATC
ACTCTTACCCCTGTTCTTATCAGACCTGGTGAGCACCTTGTTATCGATAGAATCATCCTTCAAGAATG
ACTGTGGCACGATAGCATCAACATCGTAATCAGAGAGCCTGTTGATATCCAACTCTTGATCCACATAC
ATATCCCTTCCGTTCTGGAGGTAGTAGAGGTAGAGCTTCTCATTCTGGAGCTGAGTGTTCTCAACAGG
GTGCTCTTTGAGGATCTGAGATCCAAGCTCTTTGATACCTTCCTCGATCCTCTTCATCCTTTCCCTAG
AGTTCTTCTGTCCCTTCTGAGTGGTCTGGTTCTCTCTAGCCATTTCGATCACGATGTTCTCAGGCTTA
TGCCTTCCCATCACCTTCACCAACTCATCCACAACCTTCACAGTCTGGAGGATTCCCTTCTTGATTGC
AGGAGATCCAGCGAGGTTAGCGATATGCTCATGGAGACTATCACCCTGTCCTGAAACCTGAGCCTTCT
GGATATCCTCTTTAAAGGTGAGAGAATCATCGTGGATGAGCTGCATGAAGTTTCTGTTAGCGAATCCA

TCAGACTTGAGGAAATCAAGGATTGTCTTTCCAGACTGCTTATCCCTGATTCCGTTAATGAGCTTTCT
TGAGAGCCTTCCCCAACCAGTGTATCTTCTTCTCTTCAACTGCTTCATCACCTTATCATCGAAGAGAT
GAGCGTAGGTCTTGAGCCTTTCTTCAATCATCTCTCTATCTTCAAAGAGGGTGAGGGTAAGAACGATA
TCCTCCAAGATATCCTCGTTTTCCTCGTTATCCAAGAAATCCTTATCCTTAATGATCTTGAGGAGATC
GTGGTAGGTTCCGAGAGATGCGTTGAACCTATCCTCAACACCAGAAATCTCAACTGAATCGAAGCACT
CGATTTTCTTGAAGTAATCCTCTTTGAGCTGCTTCACGGTCACCTTTCTGTTGGTCTTGAACAAGAGA
TCAACGATAGCCTTCTTTTGCTCACCTGACAAAAAAGCAGGCTTCCTCATTCCCTCGGTCACGTACTT
AACCTTGGTCAACTCGTTGTACACGGTGAAGTACTCGTAGAGCAAAGAGTGCTTAGGGAGCACCTTCT
CGTTTGGAAGGTTCTTATCGAAGTTGGTCATCCTCTCGATGAAAGACTGAGCACTAGCACCCTTATCC
ACCACCTCTTCGAAGTTCCAAGGGGTGATGGTTTCCTCAGACTTTCTGGTCATCCAAGCGAATCTTGA
GTTTCCTCTAGCGAGAGGTCCCACGTAGTAAGGGATTCTGAAGGTGAGAATCTTCTCAATCTTTTCCC
TGTTATCCTTGAGGAATGGGTAGAAATCCTCTTGCCTTCTAAGGATAGCGTGCAACTCTCCGAGGTGG
ATCTGATGAGGGATAGATCCGTTATCGAAGGTCCTCTGCTTTCTGAGAAGATCCTCTCTATTGAGCTT
CACGAGGAGTTCCTCGGTTCCATCCATCTTCTCGAGGATAGGCTTGATGAACTTGTAGAACTCTTCTT
GAGATGCACCACCATCGATGTAACCAGCGTATCCGTTCTTAGACTGATCGAAGAAAATCTCTTTGTAC
TTCTCTGGGAGCTGCTGTCTAACAAGAGCCTTGAGAAGTGTGAGATCCTGGTGGTGCTCATCGTATCT
CTTGATCATAGAAGCTGAGAGTGGAGCCTTGGTGATCTCGGTGTTCACTCTGAGGATATCACTGAGGA
GGATAGCATCAGAGAGGTTCTTAGCAGCGAGGAACAAATCAGCGTACTGATCTCCGATCTGAGCGAGG
AGGTTATCGAGATCATCATCGTAGGTATCCTTTGAGAGCTGGAGCTTTGCATCCTCAGCGAGATCGAA
GTTAGACTTGAAGTTAGGGGTGAGTCCGAGAGAGAGAGCGATCAAGTTTCCGAAAAGTCCGTTCTTCT
TCTCACCAGGGAGCTGAGCAATGAGGTTCTCAAGCCTTCTTGACTTAGAGAGCCTAGCAGAGAGGATA
GCCTTAGCATCCACACCTGAAGCGTTGATAGGGTTCTCTTCGAAAAGCTGGTTGTAGGTCTGCACGAG
CTGGATGAACAACTTATCCACATCAGAGTTATCAGGGTTGAGATCACCCTCGATGAGGAAGTGTCCTC
TGAACTTGATCATGTGAGCGAGAGCGAGGTAGATGAGCCTGAGATCAGCCTTATCAGTAGAATCAACG
AGCTTCTTTCTGAGGTGGTAGATAGTAGGGTACTTCTCGTGGTATGCCACCTCATCAACGATGTTTCC
GAAGATAGGGTGCCTCTCGTGCTTCTTATCTTCTTCCACGAGGAATGACTCTTCGAGCCTGTGGAAGA
ATGAATCATCCACTTTAGCCATCTCGTTAGAGAAGATCTCTTGGAGGTAGCAGATCCTGTTCTTTCTT
CTGGTGTACCTTCTTCTAGCGGTTCTCTTGAGTCTGGTAGCCTCAGCAGTTTCACCAGAATCGAAGAG
GAGAGCACCGATAAGGTTTTTCTTGATAGAGTGCCTATCGGTGTTTCCGAGAACCTTGAACTTCTTAG
ATGGCACCTTGTACTCATCGGTGATCACAGCCCATCCCACAGAGTTAGTTCCGATATCGAGTCCGATA
GAGTACTTCTTATCAACCTTTCTCTTCTTCTTAGGAGATTCAAATTCAGATCCATCAGCAGTTCTCTT
CATCATTcaacctttctgcaggcacatcaatactgttgaggcaacacccatagaagcacggaaccaag
gagcaccgacgtccatagcacacagccaacctcgctgaccttcgactcctcttcaagtcaaggataga
caaatctatcatgcaggaaacttacgaggcacctccgcaccagtgaccgaactgaacaccttaccgag
ataacaggactgaccaaagtgggatgtacttggctaaaacgaacctaaaaaatcactcgctcgaggcc
ccaatcgacaccacgtaaaagctccagtcgggtttcaggtctagacaatcacaatatcgagcaatcgt
aacgacgtacagtgacgagggcacaaggcgcataccagtagacaagatgcagctcttcgatgtcgatg
ctctccacgcaagagagaaagaagtgaaaggaactctagggttgcggcgatcgattttaagccagacg
ctccactgcctcccagcgagctttgagagcaagcgcgaccacagcactagacgccaaagtagaagaaa
ttagggaatgattagcgaaattgcggagggttttgatataattaaatacctaggtctgccccgtttat
aataacgagcgttcgataaggactttgcatcggctccgttcaagttattgcgacttagggtcggcggc
cggccattcgcagataggcggccaattcagcatgcgagggcaagttgggaggccgacgacgaagggtt
aggaatattcaaagcctcttttccatttatatattgccctgtccatgccccttcggactctttgtcgt
gactgccggccagcatcgtcatccgaattgctcacactcctccgtctgtcacatcgtccgtgcttagg
gccggcacgccctttcaatttaacattccaaaataagatgacgtaggtttagcgaacccgactgctga
taccacaccgattgctccggtttttcgagtaccttttccgcgcttattacaagctgcgagcaaacttt
ctagaaagtagagaggtggccgaagtcgcttttggcctcccgcgcctttctccttctgcacatacttc
caaacggtggactagcatcccatagatccatgacatgcacgcaatcacatgagattttactgttgctt
tcttcttctggcctcaccgtcctccccgtcctcccttcgctttttgggacgcgggaaaacctagaagg
cttctgattgaagtttccttttctgcttgtcggcctttacactttggagaacgaagttctaatctctc
ttgaagttgaccaaatattagtatagtgaactgatcgagaagatgcgggtccgatccaataggagaga
acaaaatattacgagacagaaagagataattaagcaatttatatgcactttaaaatggaatcttttcg

attatcatcaatgagcattatattgtgaaacggatttaaaggattagctattttgaaaggtttttta
atctgtttttaagatctttagcaaattcttttaagttacccaaatatcgatacccgttaacaatca
aaaatcgatgacagagttgtgaatcatcatcaagatctgcaaatttgatatgattttgaaaatttgag
atacaaatattgttaaaataatgcatttaactgggaaatctatcaacttaccttacaattgagttatt
ttgatatgtctctcaacaatgtgtgtgactcatttgctccactagaattcgagctcagcgaaaaaaag
caccgactcggtgccactttttcaagttgataacggactagccttattttaacttgctatttctagct
ctaaaacagacataaaaaacaaaaaaaTTTCTAGTTGGTTTAACGCGTAACTAGATAGAACCGCGTCA
AGAGAGAGAGGTACCAATCATCCTTCTGAGTACTTGACGCACCGACTCGGTGCCACTTTTTCAAGTTG
ATAACGGACTAGCCTTATTTTAACTTGCTATTTCTAGCTCTAAAACAGAAGGATGATTGGTACCGAcT
ATGgcaaggtgcaatttacggatttatacactacatttgcattgtgcgttttgcttatagttttactc
tcgataagcaaaccacgtggtcccttctccgtcattgctgtcgtgaatttctgaagtaaccggaaaat
gtttcttggccgtcatgagttcaatcctgctgtgcgatcgcgttctggctcggtccaataccactccg
tctgctcgtttcggtatatatatcggacctagaggagaatggagatcgtcaagacatcccaccatgtc
cacTTACGAGGATGCACATGTGACCGAGGGACACGAAGTGATCCGTTTAAACTATCAGTGTTTGACAG
GATATATTGGCGGGTAAACCTAAGAGAAAAGAGCGTTTATTAGAATAATCGGATATTTAAAAGGGCGT
GAAAAGGTTTATCCGTTCGTCCATTTGTATGTGCATGCCAACCACAGGGTTCCCCTCGGGAGTCAGCC
GTGCGGCTGCATGAAATCCTGGCCGGTTTGTCTGATGCCAAGCTGGCGGCCTGGCCGGCCAGCTTGGC
CGCTGAAGAAACCGAGCGCCGCCGTCTAAAAAGGTGATGTGTATTTGAGTAAAACAGCTTGCGTCATG
CGGTCGCTGCGTATATGATGCGATGAGTAAATAAACAAATACGCAAGGGGAACGCATGAAGGTTATCG
CTGTACTTAACCAGAAAGGCGGGTCAGGCAAGACGACCATCGCAACCCATCTAGCCCGCGCCCTGCAA
CTCGCCGGGGCCGATGTTCTGTTAGTCGATTCCGATCCCCAGGGCAGTGCCCGCGATTGGGCGGCCGT
GCGGGAAGATCAACCGCTAACCGTTGTCGGCATCGACCGCCCGACGATTGACCGCGACGTGAAGGCCA
TCGGCCGGCGCGACTTCGTAGTGATCGACGGAGCGCCCCAGGCGGCGGACTTGGCTGTGTCCGCGATC
AAGGCAGCCGACTTCGTGCTGATTCCGGTGCAGCCAAGCCCTTACGACATATGGGCCACCGCCGACCT
GGTGGAGCTGGTTAAGCAGCGCATTGAGGTCACGGATGGAAGGCTACAAGCGGCCTTTGTCGTGTCGC
GGGCGATCAAAGGCACGCGCATCGGCGGTGAGGTTGCCGAGGCGCTGGCCGGGTACGAGCTGCCCATT
CTTGAGTCCCGTATCACGCAGCGCGTGAGCTACCCAGGCACTGCCGCCGCCGGCACAACCGTTCTTGA
ATCAGAACCCGAGGGCGACGCTGCCCGCGAGGTCCAGGCGCTGGCCGCTGAAATTAAATCAAAACTCA
TTTGAGTTAATGAGGTAAAGAGAAAATGAGCAAAAGCACAAACACGCTAAGTGCCGGCCGTCCGAGCG
CACGCAGCAGCAAGGCTGCAACGTTGGCCAGCCTGGCAGACACGCCAGCCATGAAGCGGGTCAACTTT
CAGTTGCCGGCGGAGGATCACACCAAGCTGAAGATGTACGCGGTACGCCAAGGCAAGACCATTACCGA
GCTGCTATCTGAATACATCGCGCAGCTACCAGAGTAAATGAGCAAATGAATAAATGAGTAGATGAATT
TTAGCGGCTAAAGGAGGCGGCATGGAAAATCAAGAACAACCAGGCACCGACGCCGTGGAATGCCCCAT
GTGTGGAGGAACGGGCGGTTGGCCAGGCGTAAGCGGCTGGGTTGTCTGCCGGCCCTGCAATGGCACTG
GAACCCCCAAGCCCGAGGAATCGGCGTGACGGTCGCAAACCATCCGGCCCGGTACAAATCGGCGCGGC
GCTGGGTGATGACCTGGTGGAGAAGTTGAAGGCCGCGCAGGCCGCCCAGCGGCAACGCATCGAGGCAG
AAGCACGCCCCGGTGAATCGTGGCAAGCGGCCGCTGATCGAATCCGCAAAGAATCCCGGCAACCGCCG
GCAGCCGGTGCGCCGTCGATTAGGAAGCCGCCCAAGGGCGACGAGCAACCAGATTTTTTCGTTCCGAT
GCTCTATGACGTGGGCACCCGCGATAGTCGCAGCATCATGGACGTGGCCGTTTTCCGTCTGTCGAAGC
GTGACCGACGAGCTGGCGAGGTGATCCGCTACGAGCTTCCAGACGGGCACGTAGAGGTTTCCGCAGGG
CCGGCCGGCATGGCCAGTGTGTGGGATTACGACCTGGTACTGATGGCGGTTTCCCATCTAACCGAATC
CATGAACCGATACCGGGAAGGGAAGGGAGACAAGCCCGGCCGCGTGTTCCGTCCACACGTTGCGGACG
TACTCAAGTTCTGCCGGCGAGCCGATGGCGGAAAGCAGAAAGACGACCTGGTAGAAACCTGCATTCGG
TTAAACACCACGCACGTTGCCATGCAGCGTACGAAGAAGGCCAAGAACGGCCGCCTGGTGACGGTATC
CGAGGGTGAAGCCTTGATTAGCCGCTACAAGATCGTAAAGAGCGAAACCGGGCGGCCGGAGTACATCG
AGATCGAGCTAGCTGATTGGATGTACCGCGAGATCACAGAAGGCAAGAACCCGGACGTGCTGACGGTT
CACCCCGATTACTTTTTGATCGATCCCGGCATCGGCCGTTTTCTCTACCGCCTGGCACGCCGCGCCGC
AGGCAAGGCAGAAGCCAGATGGTTGTTCAAGACGATCTACGAACGCAGTGGCAGCGCCGGAGAGTTCA
AGAAGTTCTGTTTCACCGTGCGCAAGCTGATCGGGTCAAATGACCTGCCGGAGTACGATTTGAAGGAG
GAGGCGGGGCAGGCTGGCCCGATCCTAGTCATGCGCTACCGCAACCTGATCGAGGGCGAAGCATCCGC
CGGTTCCTAATGTACGGAGCAGATGCTAGGGCAAATTGCCCTAGCAGGGGAAAAAGGTCGAAAAAGCT
TCTTTCCTGTGGATAGCACGTACATTGGGAACCCAAAGCCGTACATTGGGAACCGGAACCCGTACATT

GGGAACCCAAAGCCGTACATTGGGAACCGGTCACACATGTAAGTGACTGATATAAAAGAGAAAAAAGG
CGATTTTTCCGCCTAAAACTCTTTAAAACTTATTAAAACTCTTAAAACCCGCCTGGCCTGTGCATAAC
TGTCTGGCCAGCGCACAGCCGAACAGCTGCAAAAAGCGCCTACCCTTCGGTCGCTGCGCTCCCTACGC
CCCGCCGCTTCGCGTCGGCCTATCGCGGCCGCTGGCCGCTCAAAAATGGCTGGCCTACGGCCAGGCAA
TCTACCAGGGCGCGGACAAGCCGCGCCGTCGCCACTCGACCGCCGGCGCCCACATCAAGGCTCCGAGT
GCGCGGAACCCCTATTTGTTTATTTTTCTAAATACATTCAAATATGTATCCGCTCATGAGACAATAAC
CCTGATAAATGCTTCAATAATATTGAAAAAGGAAGAGTATGGCTAAAATGAGAATATCACCGGAATTG
AAAAAACTGATCGAAAAATACCGCTGCGTAAAAGATACGGAAGGAATGTCTCCTGCTAAGGTATATAA
GCTGGTGGGAGAAAATGAAAACCTATATTTAAAAATGACGGACAGCCGGTATAAAGGGACCACCTATG
ATGTGGAACGGGAAAAGGACATGATGCTATGGCTGGAAGGAAAGCTGCCTGTTCCAAAGGTCCTGCAC
TTTGAACGGCATGATGGCTGGAGCAATCTGCTCATGAGTGAGGCCGATGGCGTCCTTTGCTCGGAAGA
GTATGAAGATGAACAAAGCCCTGAAAAGATTATCGAGCTGTATGCGGAGTGCATCAGGCTCTTTCACT
CCATCGACATATCGGATTGTCCCTATACGAATAGCTTAGACAGCCGCTTAGCCGAATTGGATTACTTA
CTGAATAACGATCTGGCCGATGTGGATTGCGAAAACTGGGAAGAGGACACTCCATTTAAAGATCCGCG
CGAGCTGTATGATTTTTTAAAGACGGAAAAGCCCGAAGAGGAACTTGTCTTTTCCCACGGCGACCTGG
GAGACAGCAACATCTTTGTGAAAGATGGCAAAGTAAGTGGCTTTATTGATCTTGGGAGAAGCGGCAGG
GCGGACAAGTGGTATGACATTGCCTTCTGCGTCCGGTCGCTCAGGGAGGATATCGGGGAAGAACAGTA
TGTCGAGCTATTTTTTGACTTACTGGGGATCAAGCCTGATTGGGAGAAAATAAAATATTATATTTTAC
TGGATGAATTGTTTTAGCTGTCAGACCAAGTTTACTCATATATACTTTAGATTGATTTAAAACTTCAT
TTTTAATTTAAAAGGATCTAGGTGAAGATCCTTTTTGATAATCTCATGACCAAAATCCCTTAACGTGA
GTTTTCGTTCCACTGAGCGTCAGACCCCGTAGAAAAGATCAAAGGATCTTCTTGAGATCCTTTTTTTC
TGCGCGTAATCTGCTGCTTGCAAACAAAAAAACCACCGCTACCAGCGGTGGTTTGTTTGCCGGATCAA
GAGCTACCAACTCTTTTTCCGAAGGTAACTGGCTTCAGCAGAGCGCAGATACCAAATACTGTTCTTCT
AGTGTAGCCGTAGTTAGGCCACCACTTCAAGAACTCTGTAGCACCGCCTACATACCTCGCTCTGCTAA
TCCTGTTACCAGTGGCTGCTGCCAGTGGCGATAAGTCGTGTCTTACCGGGTTGGACTCAAGACGATAG
TTACCGGATAAGGCGCAGCGGTCGGGCTGAACGGGGGGTTCGTGCACACAGCCCAGCTTGGAGCGAAC
GACCTACACCGAACTGAGATACCTACAGCGTGAGCTATGAGAAAGCGCCACGCTTCCCGAAGGGAGAA
AGGCGGACAGGTATCCGGTAAGCGGCAGGGTCGGAACAGGAGAGCGCACGAGGGAGCTTCCAGGGGGA
AACGCCTGGTATCTTTATAGTCCTGTCGGGTTTCGCCACCTCTGACTTGAGCGTCGATTTTTGTGATG
CTCGTCAGGGGGGCGGAGCCTATGGAAAAACGCCAGCAACGCGGCCTTTTTACGGTTCCTGCTCGGAT
CTGTTGGACCGGACAGTAGTCATGGTTGATGGGCTGCCTGTATCGAGTGGTGATTTTGTGCCGAGCTG
CCGGTCGGGGAGCTGTTGGCTGGCTGGTGGCAGGATATATTGTGGTGTAAACAAATTGACGCTTAGAC
AACTTAATAACACATTGCGGACGTTTTTAATGTACTGGGGTTGAACACTCTGtgccgaattcggatcc
agcgtcgatctagtaacatagatgacaccgcgcgcgataatttatcctagtttgcgcgctatatttg
ttttctatcgcgtattaaatgtataattgcgggactctaatcataaaaacccatctcataaataacgt
catgcattacatgttaattattacatgcttaacgtaattcaacagaaattatatgataatcatcgcaa
gaccggcaacaggattcaatcttaagaaactttattgccaaatgtttgaacgatctgcttgacaagcc
tattcctttgccctcggacgagtgctggggcgtcggtttccactatcggcgagtacttctacacagcc
atcggtccagacggccgcgcttctgcgggcgatttgtgtacgcccgacagtcccggctccggatcgga
cgattgcgtcgcatcgaccctgcgcccaagctgcatcatcgaaattgccgtcaaccaagctctgatag
agttggtcaagaccaatgcggagcatatacgcccggagccgcggcgatcctgcaagctccggatgcct
ccgctcgaagtagcgcgtctgctgctccatacaagccaaccacggcctccagaagaagatgttggcga
cctcgtattgggaatccccgaacatcgcctcgctccagtcaat

>2_TYPEX_007
GAACACTCTGtgccgaattcggatccagcgtcgatctagtaacatagatgacaccgcgcgcgataatt
tatcctagtttgcgcgctatatttgttttctatcgcgtattaaatgtataattgcgggactctaatc
ataaaaacccatctcataaataacgtcatgcattacatgttaattattacatgcttaacgtaattcaa
cagaaattatatgataatcatcgcaagaccggcaacaggattcaatcttaagaaactttattgccaaa
tgtttgaacgatctgcttgacaagcctattcctttgccctcggacgagtgctggggcgtcggtttcca
ctatcggcgagtacttctacacagccatcggtccagacggccgcgcttctgcgggcgatttgtgtacg

cccgacagtcccggctccggatcggacgattgcgtcgcatcgaccctgcgcccaagctgcatcatcga
aattgccgtcaaccaagctctgatagagttggtcaagaccaatgcggagcatatacgcccggagccgc
ggcgatcctgcaagctccggatgcctccgctcgaagtagcgcgtctgctgctccatacaagccaacca
cggcctccagaagaagatgttggcgacctcgtattgggaatccccgaacatcgcctcgctccagtcaa
tgaccgctgttatgcggccattgtccgtcaggacattgttggagccgaaatccgcgtgcacgagatgc
cggacttcggggcagtcctcggcccaaagcatcagctcatcgagagcctgcgcgacggacgcactgac
ggtgtcgtccatcacagtttgccagtgatacacatggggatcagcaatcgcgcatatgaaatcacgcc
atgtagtgtattgaccgattccttgcggtccgaatgggccgaacccgctcgtctggctaagatcggcc
gcagcgatcgcatccatggcctccgcgaccggctgcagttatcatcatcatagacacacgaaata
aagtaatcagattatcagttaaagctatgtaatatttacaccataaccaatcaattaaaaaatagatc
agtttaaagaaagatcaaagctcaaaaaaataaaaagagaaaagggtcctaaccaagaaaatgaagga
gaaaaactagaaatttacctgcagaacagcgggcagttcggtttcaggcaggtcttgcaacgtgacac
cctgggcacggcgggagatgcaataggtcaggctctcgctgaattccccaatgtcaagcacttccgga
atcgggagcgcggccgatgcaaagtgccgataaacataacgatctttgtagaaaccatcggcgcagct
atttacccgcaggacatatccacgccctcctacatcgaagctgaaagcacgagattcttcgccctccg
agagctgcatcaggtcggacacgctgtcgaacttttcgatcagaaacttctcgacagacgtcgcggtg
agttcaggctttttcattgcgtgtcctctccaaatgaaatgaacttccttatatagaggaagggtctt
gcgaaggatagtgggattgtgcgtcatcccttacgtcagtggagatgtcacatcaatccacttgcttt
gtagacgtggttggaacctcttctttttccacgatgctcctcgtgggtgggggtccatctttgggacc
actgtcggcagagagatcttgaatgatagcctttcctttatcgcaatgatggcatttgtaggagccac
cttccttttctactgtcctttcgatgaagtgacagatagctgggcaatggaatccgaggaggtttccc
gaaattatcctttgttgaaaagtctcaatagccctttgatcttctgagactgtatctttgacattttt
ggagtagaccagagtgtcgtgctccaccatgttgacctccGCAAGAATTCAAGCTTAGCGatctggat
tttagtactggattttggttttaggaattagaaattttattgatagaagtattttacaaatacaaata
catactaagggtttcttatatgctcaacacatgagcgaaaccctataggaaccctaattcccttatct
gggaactactcacacattattatggagaaactcgagcttgtcgatcgactctagctagagaagcCTAG
AGCGGTTGATCTGTGAGGTCTGGCCTTGTGCCATGCGCTTCCGCGAGAATATCCAGACAGTTGTGTTG
GAGGCCCTCCTCTGGGAGTGGCAGCAGTGTCGCCGGATTGAGGGCGACCACTGGGCCGAACTGGACGC
GATCGGTGTCGAGGAGGAGCGCTTGGTAATGGGTCATGCGCGCATTGGAGAGCCACCTATCCGGCGGC
TGTTTGACGAGGGCTTCCACGGCATGTGGCGCGAGGATCACCAGCGGCTGGCCCATTGTGAGTTTGCC
GGCATCCTTTGTGAGCACGGCAATCGCGGCGACCATGCGCAGACACGGTGGCCAGCCGGCGGCCACTG
GATCGAGTTTTTTGGAGAGATAGGCCACTGGCCTCCTCCACGGGCCGAGCTTCTGTGTGAGCACGCCC
TTCGCGTAGCCTTGCTTCTCATCGACGAACAGCTCGAACGGCTTTGTGAGATCTGGGAGCCCGAGCGC
TGGGGCGGTGAGGAGCGCTTGTTTGATCTCTTGGTAGGCTTTCTGCTGATCCGGCCCCCAGTTAAACA
GTGTGCCCGGCTTGGTGAGTGGGTAGAGTGGCGCGGCCATCTCGGCAAAGCCCGGGATGAAGAGGCGA
CAGAAGCCGGCTTTGCCGAGAAACTCGCGCAGCTGGCGTGGTGTTTTCGGTGTCGGTTGCCCCATCAC
TGTTTCTTTGCGCGCTTCGGTGAGCCAGCGTTGGCCTTCCTTGAGCAGATACCCGAGGTACTTCACTT
GCTTCTGGCAGATTTGGGCCTTTTTCGCGGACGCGCGATAGCCGAGGTTCCCGAGTGTTTGGAGGAGC
GCGCGTGTGCCTTGTTGACAATCCAGCTCGGAGGTCGCGGCGAGGAGGAGATCATCCACATACTGGAG
GAGGATGAGGTCCGGATGCTGGATGCGAAAATCCGCGAGGTCCCTATGGAGCGCCTCGTTGAAGAGTG
TCGGGCTGTTCTTGAAGCCTTGCGGGAGGCGTGTCCATGTCAGTTGGCCGGAGATGCCCATCTCCGGG
TCGCGCCACTCGAAGGCGAAGAGCGGCTGGCTTGTTGGGTGGAGGCGGAGACAGAAGAAGGCGTCTTT
GAGGTCCAGCACTGTATACCACTGGTGGGATGGTGGGAGCCCGGAGAGCAGATTATACGGGTTCGGGA
CGGTCGGGTGGATaTCTTCCACGCGCTTGTTGACCTCGCGGAGATCTTGGACCGGGCGATAGTCGTTG
GTGCCCGGCTTTTTGACCGGGAGGAGTGGTGTATTCCACGGGGATTGGCACGGCACGAGAATGCCTTG
ATCGAGGAGGCGCTGAATGTGCGGCTTAATCCCGAGCCTCGCCTCTTGGCTCATCGGGTACTGCTTGA
TGGACACCGGTGTGGAGGTCGCCTTCAGCGGGATAATGAGTGGGGCTTGGCGGACGGCGAGGCCCATG
CCGCCGGTTTCGGCCCACGCTTGTGGGAAGTCGCTGAGCCATGTGCTGCCGAGGGACACGTCCGGTTC
TTTGGAGGTTTCGTGGAGGCGATACTCATCCTCGATGTTGAGGGTAGGCCTAGATCCTCCGGAAGAGC
CTCCGCTAGATTCTGGTGTAGCGCTTTCGGATGTTCCTGGTGTCTCGGAGCCGCTAGATCCTCCGGAA
GATCCTCCGCTGAGCTCCACCTTCCGCTTCTTCTTTGGCGAACCAAGGAGGGATGTTTGTGGCCTTGG
GCCGCGTGGGCCGCGCGGCTTTTTCGGGCAGTCTTTGGCCCAGTGGCCCTTCTCCTTGCAGTAGGCAC

ACTGATCCCTATCGAGCTGGGACCTTCTGCGTTCTCCGCCCTGGCGGTCCTGCTTTTGGCCGGACACC
ACTGTGGCAGATTCTGGGGTGGCCGACTCGGAGGTGCCTGGCGTTTCGCTCCCGGAACCTGAATCACC
ACCGAGCTGTGAGAGATCGATCCTAGTCTCGTAGAGTCCAGTGATAGACTGATGGATGAGGGTAGCAT
CGAGCACTTCTTTGGTAGAGGTGTATCTCTTCCTATCGATGGTTGTATCGAAGTACTTGAAAGCAGCA
GGAGCACCGAGGTTGGTAAGGGTGAAGAGATGGATGATGTTCTCTGCCTGTTCCCTGATAGGCTTATC
TCTGTGCTTGTTGTAAGCAGACAACACCTTATCGAGGTTTGCATCAGCGAGGATCACCCTTTTAGAGA
ACTCAGAGATCTGCTCGATGATCTCATCCAAGTAGTGCTTGTGCTGCTCAACGAAAAGTTGCTTCTGC
TCGTTATCTTCTGGAGATCCCTTCAACTTCTCGTAGTGAGAAGCGAGGTAAAGAAAGTTAACGTACTT
AGATGGGAGAGCAAGCTCGTTTCCCTTTTGAAGCTCACCAGCAGAAGCGAGCATCCTCTTTCTACCGT
TCTCGAGTTCGAAGAGTGAGTACTTTGGGAGCTTGATGATGAGATCCTTCTTAACCTCTTTGTATCCC
TTAGCCTCGAGGAAATCGATTGGGTTCTTCTCGAAAGATGACCTTTCCATGATAGTGATTCCGAGAAG
TTCCTTAACAGACTTGAGCTTCTTACTCTTTCCCTTCTCAACCTTAGCCACAACGAGAACAGAGTAAG
CCACGGTAGGAGAATCGAAACCACCGTATTTCTTAGGGTCCCAATCCTTCTTCCTAGCAATGAGCTTA
TCAGAGTTCCTCTTAGGGAGGATAGACTCTTTAGAGAATCCACCGGTCTGCACCTCGGTTTTCTTAAC
GATGTTCACCTGTGGCATAGAGAGCACCTTTCTAACGGTAGCGAAATCCCTTCCCTTATCCCACACGA
TCTCACCTGTTTCACCGTTTGTCTCGATGAGTGGCCTCTTTCTGATCTCACCGTTAGCGAGGGTAATC
TCGGTCTTGAAGAAATTCATGATGTTAGAGTAGAAGAAATACTTAGCGGTAGCCTTTCCGATCTCTTG
CTCAGACTTAGCGATCATCTTCCTCACATCGTACACCTTGTAATCACCGTACACGAACTCTGACTCGA
GCTTAGGATACTTCTTGATGAGAGCGGTTCCAACAACAGCGTTAAGGTAAGCATCGTGAGCGTGGTGG
TAGTTGTTGATTTCCCTCACCTTGTAGAATTGGAAATCCTTTCTGAAATCAGACACGAGCTTTGACTT
GAGGGTGATAACCTTCACTTCCCTGATCAACTTATCGTTCTCATCGTACTTGGTGTTCATCCTAGAAT
CGAGGATCTGTGCAACGTGCTTAGTGATCTGCCTGGTTTCCACAAGCTGCCTCTTGATGAATCCTGCC
TTATCCAATTCAGAGAGTCCTCCCCTCTCAGCCTTAGTCAAGTTATCGAACTTTCTCTGAGTGATGAG
CTTAGCGTTGAGGAGCTGCCTCCAATAGTTCTTCATTTTCTTCACAACCTCTTCACTTGGCACGTTAT
CACTCTTACCCCTGTTCTTATCAGACCTGGTGAGCACCTTGTTATCGATAGAATCATCCTTCAAGAAT
GACTGTGGCACGATAGCATCAACATCGTAATCAGAGAGCCTGTTGATATCCAACTCTTGATCCACATA
CATATCCCTTCCGTTCTGGAGGTAGTAGAGGTAGAGCTTCTCATTCTGGAGCTGAGTGTTCTCAACAG
GGTGCTCTTTGAGGATCTGAGATCCAAGCTCTTTGATACCTTCCTCGATCCTCTTCATCCTTTCCCTA
GAGTTCTTCTGTCCCTTCTGAGTGGTCTGGTTCTCTCTAGCCATTTCGATCACGATGTTCTCAGGCTT
ATGCCTTCCCATCACCTTCACCAACTCATCCACAACCTTCACAGTCTGGAGGATTCCCTTCTTGATTG
CAGGAGATCCAGCGAGGTTAGCGATATGCTCATGGAGACTATCACCCTGTCCTGAAACCTGAGCCTTC
TGGATATCCTCTTTAAAGGTGAGAGAATCATCGTGGATGAGCTGCATGAAGTTTCTGTTAGCGAATCC
ATCAGACTTGAGGAAATCAAGGATTGTCTTTCCAGACTGCTTATCCCTGATTCCGTTAATGAGCTTTC
TTGAGAGCCTTCCCCAACCAGTGTATCTTCTTCTCTTCAACTGCTTCATCACCTTATCATCGAAGAGA
TGAGCGTAGGTCTTGAGCCTTTCTTCAATCATCTCTCTATCTTCAAAGAGGGTGAGGGTAAGAACGAT
ATCCTCCAAGATATCCTCGTTTTCCTCGTTATCCAAGAAATCCTTATCCTTAATGATCTTGAGGAGAT
CGTGGTAGGTTCCGAGAGATGCGTTGAACCTATCCTCAACACCAGAAATCTCAACTGAATCGAAGCAC
TCGATTTTCTTGAAGTAATCCTCTTTGAGCTGCTTCACGGTCACCTTTCTGTTGGTCTTGAACAAGAG
ATCAACGATAGCCTTCTTTTGCTCACCTGACAAAAAAGCAGGCTTCCTCATTCCCTCGGTCACGTACT
TAACCTTGGTCAACTCGTTGTACACGGTGAAGTACTCGTAGAGCAAAGAGTGCTTAGGGAGCACCTTC
TCGTTTGGAAGGTTCTTATCGAAGTTGGTCATCCTCTCGATGAAAGACTGAGCACTAGCACCCTTATC
CACCACCTCTTCGAAGTTCCAAGGGGTGATGGTTTCCTCAGACTTTCTGGTCATCCAAGCGAATCTTG
AGTTTCCTCTAGCGAGAGGTCCCACGTAGTAAGGGATTCTGAAGGTGAGAATCTTCTCAATCTTTTCC
CTGTTATCCTTGAGGAATGGGTAGAAATCCTCTTGCCTTCTAAGGATAGCGTGCAACTCTCCGAGGTG
GATCTGATGAGGGATAGATCCGTTATCGAAGGTCCTCTGCTTTCTGAGAAGATCCTCTCTATTGAGCT
TCACGAGGAGTTCCTCGGTTCCATCCATCTTCTCGAGGATAGGCTTGATGAACTTGTAGAACTCTTCT
TGAGATGCACCACCATCGATGTAACCAGCGTATCCGTTCTTAGACTGATCGAAGAAAATCTCTTTGTA
CTTCTCTGGGAGCTGCTGTCTAACAAGAGCCTTGAGAAGTGTGAGATCCTGGTGGTGCTCATCGTATC
TCTTGATCATAGAAGCTGAGAGTGGAGCCTTGGTGATCTCGGTGTTCACTCTGAGGATATCACTGAGG
AGGATAGCATCAGAGAGGTTCTTAGCAGCGAGGAACAAATCAGCGTACTGATCTCCGATCTGAGCGAG
GAGGTTATCGAGATCATCATCGTAGGTATCCTTTGAGAGCTGGAGCTTTGCATCCTCAGCGAGATCGA
AGTTAGACTTGAAGTTAGGGGTGAGTCCGAGAGAGAGAGCGATCAAGTTTCCGAAAAGTCCGTTCTTC

TTCTCACCAGGGAGCTGAGCAATGAGGTTCTCAAGCCTTCTTGACTTAGAGAGCCTAGCAGAGAGGAT
AGCCTTAGCATCCACACCTGAAGCGTTGATAGGGTTCTCTTCGAAAAGCTGGTTGTAGGTCTGCACGA
GCTGGATGAACAACTTATCCACATCAGAGTTATCAGGGTTGAGATCACCCTCGATGAGGAAGTGTCCT
CTGAACTTGATCATGTGAGCGAGAGCGAGGTAGATGAGCCTGAGATCAGCCTTATCAGTAGAATCAAC
GAGCTTCTTTCTGAGGTGGTAGATAGTAGGGTACTTCTCGTGGTATGCCACCTCATCAACGATGTTTC
CGAAGATAGGGTGCCTCTCGTGCTTCTTATCTTCTTCCACGAGGAATGACTCTTCGAGCCTGTGGAAG
AATGAATCATCCACTTTAGCCATCTCGTTAGAGAAGATCTCTTGGAGGTAGCAGATCCTGTTCTTTCT
TCTGGTGTACCTTCTTCTAGCGGTTCTCTTGAGTCTGGTAGCCTCAGCAGTTTCACCAGAATCGAAGA
GGAGAGCACCGATAAGGTTTTTCTTGATAGAGTGCCTATCGGTGTTTCCGAGAACCTTGAACTTCTTA
GATGGCACCTTGTACTCATCGGTGATCACAGCCCATCCCACAGAGTTAGTTCCGATATCGAGTCCGAT
AGAGTACTTCTTATCAACCTTTCTCTTCTTCTTAGGAGATTCAAATTCAGATCCATCAGCAGTTCTCT
TCATCATTcaacctttctgcaggcacatcaatactgttgaggcaacacccatagaagcacggaaccaa
ggagcaccgacgtccatagcacacagccaacctcgctgaccttcgactcctcttcaagtcaaggatag
acaaatctatcatgcaggaaacttacgaggcacctccgcaccagtgaccgaactgaacaccttaccga
gataacaggactgaccaaagtgggatgtacttggctaaaacgaacctaaaaatcactcgctcgaggc
cccaatcgacaccacgtaaaagctccagtcgggtttcaggtctagacaatcacaatatcgagcaatcg
taacgacgtacagtgacgagggcacaaggcgcataccagtagacaagatgcagctcttcgatgtcgat
gctctccacgcaagagagaaagaagtgaaaggaactctagggttgcggcgatcgattttaagccagac
gctccactgcctcccagcgagctttgagagcaagcgcgaccacagcactagacgccaaagtagaagaa
attagggaatgattagcgaaattgcggagggttttgatataattaaatacctaggtctgccccgttta
taataacgagcgttcgataaggactttgcatcggctccgttcaagttattgcgacttagggtcggcgg
ccggccattcgcagataggcggccaattcagcatgcgagggcaagttgggaggccgacgacgaagggt
taggaatattcaaagcctcttttccatttatatattgccctgtccatgccccttcggactctttgtcg
tgactgccggccagcatcgtcatccgaattgctcacactcctccgtctgtcacatcgtccgtgcttag
ggccggcacgccctttcaatttaacattccaaaataagatgacgtaggtttagcgaacccgactgctg
ataccacaccgattgctccggtttttcgagtacctttccgcgcttattacaagctgcgagcaaactt
tctagaaagtagagaggtggccgaagtcgcttttggcctcccgcgcctttctccttctgcacatactt
ccaaacggtggactagcatcccatagatccatgacatgcacgcaatcacatgagattttactgttgct
ttcttcttctggcctcaccgtcctccccgtcctcccttcgctttttgggacgcgggaaaacctagaag
gcttctgattgaagtttccttttctgcttgtcggcctttacactttggagaacgaagttctaatctct
cttgaagttgaccaaatattagtatagtgaactgatcgagaagatgcgggtccgatccaataggagag
aacaaaatattacgagacagaaagagataattaagcaatttatatgcactttaaaatggaatcttttc
gattatcatcaatgagcattatattgtgaaacggatttaaaggattagctattttgaaaggtttttt
aatctgtttttaagatctttagcaaattctttttaagttacccaaatatcgatacccgttaacaatc
aaaaatcgatgacagagttgtgaatcatcatcaagatctgcaaatttgatatgattttgaaaatttga
gatacaaatattgttaaaataatgcatttaactgggaaatctatcaacttaccttacaattgagttat
tttgatatgtctctcaacaatgtgtgtgactcatttgctccactagaattcgagctcagcgaaaaaa
gcaccgactcggtgccactttttcaagttgataacggactagccttattttaacttgctatttctagc
tctaaaacagacataaaaaacaaaaaaaTTTCTAGTTGGTTTAACGCGTAACTAGATAGAACCGCGTC
AAGAGAGAGAGGTACCAATCATCCTTCTGAGTACTTGACGCACCGACTCGGTGCCACTTTTTCAAGTT
GATAACGGACTAGCCTTATTTTAACTTGCTATTTCTAGCTCTAAAACAGAAGGATGATTGGTACCGAc
TATGgcaaggtgcaatttacggatttatacactacatttgcattgtgcgttttgcttatagttttact
ctcgataagcaaaccacgtggtcccttctccgtcattgctgtcgtgaatttctgaagtaaccggaaaa
tgtttcttggccgtcatgagttcaatcctgctgtgcgatcgcgttctggctcggtccaataccactcc
gtctgctcgtttcggtatatatatcggacctagaggagaatggagatcgtcaagacatcccaccatgt
ccacaaatttgtgctccatcgcgggtttctttccatcgtgaaagcgatgcatacaccaaaaaacgaag
cataactgcgcatgccccaagagcatcgtttagagcaacaggaataagacttagatttgattcgatat
cgtaaagatagcgaccatgggacacaccacttgtttctagaacttcgaaccccgggggaaaatagttg
atggggccttcccatcctttttgctatattacttggtattcccgaagaagggcttctttggacggaga
ggagtgtatgtacgatgctagctccgtgttttcgtgttttcttgtttccccaaacctgaaggatttgg
gtggaagtaccaaggagattcaactgtccagttgtgcgggctacccaaatgaaagaactcccacacgc
gcaaatttgtttcctacattgagaaataactgcaaaagcctgcacgttgctgtaagaaaaataatctg

taggaaattaggaaactacagagctcgtctgaaaattcccagctttcatttcaggtttacggtatttt
atttctgacgtgacagaaataaaattggaaattggaaattcattggatgcatggccaggtatcggtgc
agtcacatctttcgcacccgttttatcactacaacgttagtgcggtgtacatttcaatcgctgttgtt
atctgagcttatctcgccgttccatttgagtcatccgatccaacgagaacagacacgtctgacgcagt
cctttagaatccaggtcatttcgcattcttccccacaggaatcaagtctctaatcttgttccgatccg
tcagacaacacttgctccgtccgcagagccctgtgcgattgagtgccgagtgttcttggtgcttcatt
ttagtaaaactcgatcagcgatcgttcttggttctggacagatctgcaggtcacgagaacagctcatg
tcttttatccttagagaaatcggtcatatgtttgcagcttgtcagaacttttgcgactttattgatta
cgtgacatacgcggaacaagaggctacacaatcggaagggcgaatcggccggatgcactggcagtgga
ttccatcaagacaatacagtcgcttgcggaaacagcattcacatttgttgtcaatgcagctaggaaga
ggtaaattcacatcttgacatgctaaacaaatcagcacctttgaacataattccgttttgtatgtcaa
tacaaaagtgtactactaagccattgctaaaatgcataaatagcaatgctttcctctgagaacaaact
gattgccactaatttaatgtgaacatggattacaatagagcctgtagctcaaatacctgcgcatccct
acatcatgcagcaaaattggatagcatccgctagaacgagatttagggcgctggagctggactgatgg
gtccttttctacttcctgatggtggggacggaggaattacaacccaaggtgtccggccggtaggagcc
tttctgggtggtgggggagccgtggccagttcagtcacacgaagctgccggctggtccctactgccac
ctctgtcatgtccacgggaacatcgactgaagctttctcttgctttatcacgtccactttttcgtga
gtttacgtcccggaactaatgctctctctgtgaaagtcgtccccttgacctgccatacatacatctcg
actgaacaacttcagccatcactccTTACGAGGATGCACATGTGACCGAGGGACACGAAGTGATCCGT
TTAAACTATCAGTGTTTGACAGGATATATTGGCGGGTAAACCTAAGAGAAAAGAGCGTTTATTAGAAT
AATCGGATATTTAAAAGGGCGTGAAAAGGTTTATCCGTTCGTCCATTTGTATGTGCATGCCAACCACA
GGGTTCCCCTCGGGAGTCAGCCGTGCGGCTGCATGAAATCCTGGCCGGTTTGTCTGATGCCAAGCTGG
CGGCCTGGCCGGCCAGCTTGGCCGCTGAAGAAACCGAGCGCCGCCGTCTAAAAAGGTGATGTGTATTT
GAGTAAAACAGCTTGCGTCATGCGGTCGCTGCGTATATGATGCGATGAGTAAATAAACAAATACGCAA
GGGGAACGCATGAAGGTTATCGCTGTACTTAACCAGAAAGGCGGGTCAGGCAAGACGACCATCGCAAC
CCATCTAGCCCGCGCCCTGCAACTCGCCGGGGCCGATGTTCTGTTAGTCGATTCCGATCCCCAGGGCA
GTGCCCGCGATTGGGCGGCCGTGCGGGAAGATCAACCGCTAACCGTTGTCGGCATCGACCGCCCGACG
ATTGACCGCGACGTGAAGGCCATCGGCCGGCGCGACTTCGTAGTGATCGACGGAGCGCCCCAGGCGGC
GGACTTGGCTGTGTCCGCGATCAAGGCAGCCGACTTCGTGCTGATTCCGGTGCAGCCAAGCCCTTACG
ACATATGGGCCACCGCCGACCTGGTGGAGCTGGTTAAGCAGCGCATTGAGGTCACGGATGGAAGGCTA
CAAGCGGCCTTTGTCGTGTCGCGGGCGATCAAAGGCACGCGCATCGGCGGTGAGGTTGCCGAGGCGCT
GGCCGGGTACGAGCTGCCCATTCTTGAGTCCCGTATCACGCAGCGCGTGAGCTACCCAGGCACTGCCG
CCGCCGGCACAACCGTTCTTGAATCAGAACCCGAGGGCGACGCTGCCCGCGAGGTCCAGGCGCTGGCC
GCTGAAATTAAATCAAAACTCATTTGAGTTAATGAGGTAAAGAGAAAATGAGCAAAAGCACAAACACG
CTAAGTGCCGGCCGTCCGAGCGCACGCAGCAGCAAGGCTGCAACGTTGGCCAGCCTGGCAGACACGCC
AGCCATGAAGCGGGTCAACTTTCAGTTGCCGGCGGAGGATCACACCAAGCTGAAGATGTACGCGGTAC
GCCAAGGCAAGACCATTACCGAGCTGCTATCTGAATACATCGCGCAGCTACCAGAGTAAATGAGCAAA
TGAATAAATGAGTAGATGAATTTTAGCGGCTAAAGGAGGCGGCATGGAAAATCAAGAACAACCAGGCA
CCGACGCCGTGGAATGCCCCATGTGTGGAGGAACGGGCGGTTGGCCAGGCGTAAGCGGCTGGGTTGTC
TGCCGGCCCTGCAATGGCACTGGAACCCCCAAGCCCGAGGAATCGGCGTGACGGTCGCAAACCATCCG
GCCCGGTACAAATCGGCGCGGCGCTGGGTGATGACCTGGTGGAGAAGTTGAAGGCCGCGCAGGCCGCC
CAGCGGCAACGCATCGAGGCAGAAGCACGCCCCGGTGAATCGTGGCAAGCGGCCGCTGATCGAATCCG
CAAAGAATCCCGGCAACCGCCGGCAGCCGGTGCGCCGTCGATTAGGAAGCCGCCCAAGGGCGACGAGC
AACCAGATTTTTTCGTTCCGATGCTCTATGACGTGGGCACCCGCGATAGTCGCAGCATCATGGACGTG
GCCGTTTTCCGTCTGTCGAAGCGTGACCGACGAGCTGGCGAGGTGATCCGCTACGAGCTTCCAGACGG
GCACGTAGAGGTTTCCGCAGGGCCGGCCGGCATGGCCAGTGTGTGGGATTACGACCTGGTACTGATGG
CGGTTTCCCATCTAACCGAATCCATGAACCGATACCGGGAAGGGAAGGGAGACAAGCCCGGCCGCGTG
TTCCGTCCACACGTTGCGGACGTACTCAAGTTCTGCCGGCGAGCCGATGGCGGAAAGCAGAAAGACGA
CCTGGTAGAAACCTGCATTCGGTTAAACACCACGCACGTTGCCATGCAGCGTACGAAGAAGGCCAAGA
ACGGCCGCCTGGTGACGGTATCCGAGGGTGAAGCCTTGATTAGCCGCTACAAGATCGTAAAGAGCGAA
ACCGGGCGGCCGGAGTACATCGAGATCGAGCTAGCTGATTGGATGTACCGCGAGATCACAGAAGGCAA
GAACCCGGACGTGCTGACGGTTCACCCCGATTACTTTTTGATCGATCCCGGCATCGGCCGTTTTCTCT

ACCGCCTGGCACGCCGCGCCGCAGGCAAGGCAGAAGCCAGATGGTTGTTCAAGACGATCTACGAACGC
AGTGGCAGCGCCGGAGAGTTCAAGAAGTTCTGTTTCACCGTGCGCAAGCTGATCGGGTCAAATGACCT
GCCGGAGTACGATTTGAAGGAGGAGGCGGGGCAGGCTGGCCCGATCCTAGTCATGCGCTACCGCAACC
TGATCGAGGGCGAAGCATCCGCCGGTTCCTAATGTACGGAGCAGATGCTAGGGCAAATTGCCCTAGCA
GGGGAAAAAGGTCGAAAAAGCTTCTTTCCTGTGGATAGCACGTACATTGGGAACCCAAAGCCGTACAT
TGGGAACCGGAACCCGTACATTGGGAACCCAAAGCCGTACATTGGGAACCGGTCACACATGTAAGTGA
CTGATATAAAAGAGAAAAAAGGCGATTTTTCCGCCTAAAACTCTTTAAAACTTATTAAAACTCTTAAA
ACCCGCCTGGCCTGTGCATAACTGTCTGGCCAGCGCACAGCCGAACAGCTGCAAAAAGCGCCTACCCT
TCGGTCGCTGCGCTCCCTACGCCCCGCCGCTTCGCGTCGGCCTATCGCGGCCGCTGGCCGCTCAAAAA
TGGCTGGCCTACGGCCAGGCAATCTACCAGGGCGCGGACAAGCCGCGCCGTCGCCACTCGACCGCCGG
CGCCCACATCAAGGCTCCGAGTGCGCGGAACCCCTATTTGTTTATTTTTCTAAATACATTCAAATATG
TATCCGCTCATGAGACAATAACCCTGATAAATGCTTCAATAATATTGAAAAAGGAAGAGTATGGCTAA
AATGAGAATATCACCGGAATTGAAAAAACTGATCGAAAAATACCGCTGCGTAAAAGATACGGAAGGAA
TGTCTCCTGCTAAGGTATATAAGCTGGTGGGAGAAAATGAAAACCTATATTTAAAAATGACGGACAGC
CGGTATAAAGGGACCACCTATGATGTGGAACGGGAAAAGGACATGATGCTATGGCTGGAAGGAAAGCT
GCCTGTTCCAAAGGTCCTGCACTTTGAACGGCATGATGGCTGGAGCAATCTGCTCATGAGTGAGGCCG
ATGGCGTCCTTTGCTCGGAAGAGTATGAAGATGAACAAAGCCCTGAAAAGATTATCGAGCTGTATGCG
GAGTGCATCAGGCTCTTTCACTCCATCGACATATCGGATTGTCCCTATACGAATAGCTTAGACAGCCG
CTTAGCCGAATTGGATTACTTACTGAATAACGATCTGGCCGATGTGGATTGCGAAAACTGGGAAGAGG
ACACTCCATTTAAAGATCCGCGCGAGCTGTATGATTTTTTAAAGACGGAAAAGCCCGAAGAGGAACTT
GTCTTTTCCCACGGCGACCTGGGAGACAGCAACATCTTTGTGAAAGATGGCAAAGTAAGTGGCTTTAT
TGATCTTGGGAGAAGCGGCAGGGCGGACAAGTGGTATGACATTGCCTTCTGCGTCCGGTCGCTCAGGG
AGGATATCGGGGAAGAACAGTATGTCGAGCTATTTTTTGACTTACTGGGGATCAAGCCTGATTGGGAG
AAAATAAAATATTATATTTTACTGGATGAATTGTTTTAGCTGTCAGACCAAGTTTACTCATATATACT
TTAGATTGATTTAAAACTTCATTTTTAATTTAAAAGGATCTAGGTGAAGATCCTTTTTGATAATCTCA
TGACCAAAATCCCTTAACGTGAGTTTTCGTTCCACTGAGCGTCAGACCCCGTAGAAAAGATCAAAGGA
TCTTCTTGAGATCCTTTTTTTCTGCGCGTAATCTGCTGCTTGCAAACAAAAAAACCACCGCTACCAGC
GGTGGTTTGTTTGCCGGATCAAGAGCTACCAACTCTTTTTCCGAAGGTAACTGGCTTCAGCAGAGCGC
AGATACCAAATACTGTTCTTCTAGTGTAGCCGTAGTTAGGCCACCACTTCAAGAACTCTGTAGCACCG
CCTACATACCTCGCTCTGCTAATCCTGTTACCAGTGGCTGCTGCCAGTGGCGATAAGTCGTGTCTTAC
CGGGTTGGACTCAAGACGATAGTTACCGGATAAGGCGCAGCGGTCGGGCTGAACGGGGGGTTCGTGCA
CACAGCCCAGCTTGGAGCGAACGACCTACACCGAACTGAGATACCTACAGCGTGAGCTATGAGAAAGC
GCCACGCTTCCCGAAGGGAGAAAGGCGGACAGGTATCCGGTAAGCGGCAGGGTCGGAACAGGAGAGCG
CACGAGGGAGCTTCCAGGGGGAAACGCCTGGTATCTTTATAGTCCTGTCGGGTTTCGCCACCTCTGAC
TTGAGCGTCGATTTTTGTGATGCTCGTCAGGGGGGCGGAGCCTATGGAAAAACGCCAGCAACGCGGCC
TTTTTACGGTTCCTGCTCGGATCTGTTGGACCGGACAGTAGTCATGGTTGATGGGCTGCCTGTATCGA
GTGGTGATTTTGTGCCGAGCTGCCGGTCGGGGAGCTGTTGGCTGGCTGGTGGCAGGATATATTGTGGT
GTAAACAAATTGACGCTTAGACAACTTAATAACACATTGCGGACGTTTTTAATGTACTGGGGTT
